# NbCycB2 represses Nbwo activity via a negative feedback loop in the tobacco trichome developmemt

**DOI:** 10.1101/740126

**Authors:** Minliang Wu, Yuchao Cui, Li Ge, Lipeng Cui, Zhichao Xu, Hongying Zhang, Zhaojun Wang, Dan Zhou, Shuang Wu, Liang Chen, Hong Cui

## Abstract

The wo protein and its downstream gene, *SlCycB2* have been demonstrated to regulate the trichome development in tomato. It was shown that only gain-of-function mutant form of *wo, Wo^v^* (wo woolly motif mutant allele) could induce the increase of trichome density. However, it is still unclear the relationships between wo, *Wo^v^* and *SlCycB2* in trichome regulation. In this study, we demonstrated Nbwo (NbWo^v^) directly regulated the expressions *NbCycB2* by binding to the promoter of *NbCycB2* and its genomic sequences. As a feedback regulation, NbCycB2 negatively regulates the trichome formation by repressing Nbwo activity at protein level. We further found that the mutations of Nbwo woolly motif could prevent repression of NbWo^v^ by NbCycB2, which results in the significant increase of active *Nbwo* proteins, trichome density and branches. Our results revealed a novel reciprocal mechanism between *NbCycB2* and *Nbwo* during the trichome formation in *Nicotiana benthamiana*.

**Highlight:** *NbCycB2* is specifically expressed in trichomes of *Nicotiana benthamiana* and represses the Nbwo activity via a negative feedback loop in tobacco trichome developmemt.

## Introduction

Trichomes are the specialized epidermal protuberances locating on aerial parts of nearly all terrestrial plants. They can be classified into various types by cell numbers and shapes -- unicellular/multicellular, and glandular/non-glandular. In *Arabidopsis,* it has been well-elucidated that the development of trichomes (unicellular and non-glandular), is regulated by the trimeric MYB-bHLH-WDR protein activators complex (GL1 (Oppenheimer et al., 1991)-GL3/EGL3 (Payne et al., 2000)-TTG1(Walker et al., 1999)). This transcriptional complex activates the expression of the homeodomain protein GLABROUS2 (GL2) to induce the formation of trichomes (Rerie et al., 1994; Grebe, 2012). In addition, it triggers the expression of the single repeat R3 MYBs (including TRY (Schnittger et al., 1999), CPC (Wada et al., 1997), ETC1, ETC2, ETC3 (Kirik et al., 2004; Wester et al., 2009) and TCL2 (Gan et al., 2011)) which act as negative regulators of GL3 or EGL3 by forming a repressor complex (GL3/EGL3-TRY/CPC-TTG1) in trichome development (Wang et al., 2008; Wester et al., 2009). Thus the control of trichome development in *Arabidopsis* requires a regulatory loop including both activators and repressors (Grebe, 2012; Pattanaik et al., 2014).

The trichomes are multicellular glandular (GSTs) structure in approximately 30% of all vascular plants (Glas et al., 2012). Since many phytochemicals and compounds with economical values can be synthesized and secreted by multicellular GSTs (Mauricio and Rausher, 1997; Hollósy, 2002; Valkama et al., 2003; Freeman and Beattie, 2008), multicellular glandular trichomes have considerable economic pontential (Sallets et al., 2014; Huchelmann et al., 2017). However, it has been demonstrated that the networks regulating unicellular trichomes did not work in the development of multicellular trichomes (Serna and Martin, 2006; Yang et al., 2011; Kang et al., 2016; Yan et al., 2016).

In tomato, a HD-ZIP IV transcriptional factor, wo protein has been demonstrated to regulate the trichome initiation (Yang et al., 2011). This HD-ZIP IV member contains four conserved domains including homeodomain domain (HD), the leucine zipper domain (LZ), the steroidogenic acute regulatory protein-related lipid transfer (START) and the START-adjacent domain (SAD). However, overexpression of *wo* failed to induce the change of trichome density, and only ectopic expression of its gain-of-function mutant alleles, *Wo^v^*, could cause higher density of trichomes in tomato *(Solanum lycopersicum)* and tobacco *(Nicotiana tabacum)* (Yang et al., 2011; Yang et al., 2015). The *Wo^v^* allele has two point mutations at the C-terminal domain (Since this motif was conserved in the most wo alleles, we name it as woolly motif in this study). The sequence analysis revealed that wo protein is more similar to PROTODERMAL FACTOR2 (PDF2) and the PDF2 redundant protein -- ARABIDOPSIS THALIANA MERISTEM L1 (ATML1), both of which are involved in shoot epidermal cell differentiation (Abe et al., 2001; Ogawa et al., 2015), than GL2 in *Arabidopsis*.

In *Arabidopsis,* the ectopic expression of a constitutive active B-type cyclin induced mitotic divisions and resulted in the increase of multicellular trichomes (Schnittger et al., 2005). *SlCycB2,* a hypothetical B-type cyclin, was reported to directly interact with wo to promote the development of type I trichome (Yang et al., 2011; Yang et al., 2015). Its homologous protein in *Arabidopsis* (AT5G06270) was also found to interact with GL2 or co-repressor TOPLESS proteins (Wu and Citovsky, 2017b, a). However, as reported in a recent study, overexpression of *SICyc82* resulted in non-trichome phenotype, while suppression of *SICyc82* promoted trichomes formation in tomato (Gao et al., 2017). These inconsistent results raise the important questions: what is the function of *SlCycB2* in trichome formation and why the mutation of woolly motif can promote trichome formation?

Similar to tomato, trichomes in *Nicotiana benthamiana* are typically multicellular structures, and almost all of the trichomes in *N. benthamiana* are glandular (Fig. S1), making it a better system for studying glandular trichomes than tomato. In addition, the genome map of *N. benthamiana* has been constructed (Bombarely et al., 2012). Thus tobacco represents an excellent model plant to study the molecular mechanism of multicellular trichome formation (Goodin et al., 2008). In this study, we cloned the homologues of *wo* and *SlCycB2* in *N. benthamiana* (named *Nbwo* and *NbCycB2),* and constructed a two-point mutantion *Nbwo* allele, *NbWo^v^.* To investigate their biological functions in trichome development, we constructed overexpression and suppression transgenic lines of all genes. We demonstrated that Nbwo could positively regulate the expression of *NbCycB2* through targeting to the cis-element in *NbCycB2* promoter and its genomic DNA sequence. On the other hand, NbCycB2 could be a negative regulator of multicellular trichomes by directly binding and inhibiting Nbwo activity. The previous identified mutation in woolly motif (NbWo^v^) blocked the interaction between NbCycB2 and Nbwo, removing the repression of *Nbwo* by NbCycB2 and resulting in increased trichome density. Our results revealed the mechanisms of the interaction between *Nbwo* and *NbCycB2* in regulating the development of glandular trichomes.

## Materials and Methods

### Plant materials and growth conditions

Sterilized seeds of *N. benthamiana* were germinated and grown to seedlings on MS medium, which solidified with 0.8% (w/v) gellan gum under 26 °C, 14 h light/10 h dark conditions. Two-week-old plants were transferred to other sterilized bottle (for genetic transformation) or soil in pots to grow to maturity. All wild type and transgenic plants were grown in greenhouse under 26 °C, 14 h light/10 h dark condition.

### Sequence analysis

The protein sequences of the homologues of *wo* and *SlCycB2* were downloaded from NCBI database (http://www.ncbi.nlm.nih.gov/) and Sol genomic network (Fernandez-Pozo et al., 2015) (https://solgenomics.net/). In order to further analyze the grouping and relatedness, the aligned sequences were used to construct the phylogenetic trees in MEGA 5 by using the maximum-likelihood (ML) criterion with 100 bootstrap analysis. In addition, the relative conservation for each amino acid position in the protein sequences of Nbwo and NbCycB2 was evaluated via Weblogo (https://weblogo.berkeley.edu/) (Crooks et al., 2004), followed by the prediction of their conserved domains in SMART program (https://smart.embl-heidelberg.de/) (Letunic and Bork, 2018).

### RNA extraction and Real-time PCR

Total of RNA was extracted from various tissue of plants by using the Eastep^®^Super Total RNA Extraction Kit (Promega). The cDNA was synthesized from Dnase I treated total RNA using M-MLV 1st Strand Kit (Invitrogen). Real-time PCR (qRT-PCR) was determined by using SYBR Premix Ex Taq II (TaKaRa) and performed on ABI Stepone real-time PCR system (Applied Biosystems). L25 ribosomal protein (L18908) was used as an endogenous control (Schmidt and Delaney, 2010). Relative expression levels were determined as described previously (Guo et al., 2016). Primers are listed in Table S1.

### Plasmid construction and *N. benthamiana* transformation

To investigate the biological functions of *wo* and *SlCyc82* in *N. benthamiana,* the full-length coding sequences of two alleles of *Nbwo* and *NbCycB2* were amplified with Nhel and BamHI, Xbal and BgIll cloning sites respectively from the general cDNA of leaves (All the primers used in current study are provided in Table S1). In addition, to confirm the function of the tomato *Wo^v^* gene in tobacoo trichome formation (Yang et al., 2015), an allele *NbWo^v^* with two point mutations at the locus 2084 (T was replaced with G, result in ile-697 changed to Arg) and 2092 (G was replaced with T, result in Asp-700 changed to Tyr, Fig. S2c) of *Nbwo* was generated in *N. benthamiana* by using a Mutagenesis Kit (Toyobo).

To construct the overexpression (OE) lines of *Nbwo, NbWo^v^* and *NbCycB2,* these fragments were inserted into pCXSN-HA (Nbwo and NbWo^v^ fused with HA tag) and pCXSN-FLAG (NbCycB2 fused with Flag tag) vectors respectively under the control of CaMV 35S promoter (Chen et al., 2009). The suppression expression constructs of *Nbwo* and *NbCycB2* were performed by recombining with the RNAi vector pH7GWIWGII with LR Clonase II enzyme (Invitrogen).

To comprehensively understand the expression sites of *NbCycB2,* approximate 2880bp upstream promoter fragments were amplified by PCR using the primers shown in Table S1. The promoter fragments were then inserted into the corresponding site of pH2GW7 vector to create the promoter-driven GFP-GUS transformation by using *Pro-NbCycB2:* GFP-GUS gene fusion (Cui et al., 2015).

All of these constructs were transferred into *Agrobacterium tumefaciens* strain GV3101, which were used to generate transgenic lines by using *Agrobacterium-mediated* transformation. The gene expression levels in the transgenic lines were examined by real-time PCR and western blot.

### Subcellular localization and tissue distribution

The pCXDG vector (Chen et al., 2009) was used to analyse the subcellular localization of *Nbwo, NbWo^v^* and *NbCycB2,* which were fused with GFP driven by the CaMV 35S promoter (p35s:GFP-Nbwo, p35s:GFP-NbWoV, p35s:GFP-NbCycB2). Transformants carrying these constructs were obtained as described above and then transiently transformed into 4-week-old *N. benthamiana* leaves. After cultivation in lowlight conditions for 48-72 h, GFP was observed using confocal microscopy (LSM 780, Carl Zeiss, Jena, Germany) with staining in DAPI solution (1 mg/ml) for 15 min before observation.

The tissue distribution assays were performed as described in the previous study (Jefferson et al., 1987). GUS staining was repeated at least three independent transgenic lines.

### Yeast hybrid assays

Yeast one-hybrid assays were performed to test the specific function area of *NbCycB2* promoter binding Nbwo. The promoter of *NbCycB2,* separated into 5 fragments (E: −1027 ∼ −831 bp, D: −830 ∼ −631 bp, C: - 630 ∼ −411 bp, B: −410 ∼ −201 bp, A: −200 ∼ −1bp, Fig. 2a), were amplified and inserted into the pHIS 2 vector (Clontech) *(NbCycB2proE, NbCycB2proD, NbCycB2proC, NbCycB2proB, NbCycB2proA).* Further investigation of the targeted sequences in *NbCycB2* promoter were conducted by point mutations in the two L1-like boxes in the D fragment *(NbCycB2proD-m1,* mutant one L1-like box, changed 5’-GCAAATATTTACTC-3’ to 5’-GCGGGTGACTC-3’; *NbCycB2proD-m2,* mutant two L1-like boxes, changed 5’-GCAAATATTTACTC-3’ to 5’-GCGGGTGACTC-3’, and 5’-ATTTACTC-3’ to 5’-GGGACTCC-3’). To test the specific region of *Nbwo* genomic sequence binding itself, four genomic fragments of *Nbwo* (G1, −8 ∼ 251 bp, include T3 fragment; G2, 2169 ∼ 2522 bp, include T4 fragment; G3, 3485 ∼ 3780 bp, include T5 fragment; G4, 4333 ∼ 4660 bp, include T6 fragment, Fig. 7a), were amplified and inserted into the pHIS 2 vector (Clontech) *(Nbwo-G1, Nbwo-G2, Nbwo-G3, Nbwo-G4).* In addition, the CDSs of *Nbwo* and *NbWo^v^* were inserted into vectors pGADT7 containing the GAL4 activation domain (AD) (Clontech). The plasmids were co-transformed into the Y187 yeast strain, empty AD vector was provided as the negative control, and cultivated on SD/-Leu/-Trp (-L-W) medium and tested on SD/-Leu/-His/-Trp (-L-W-H) with 60 mM 3-amino-1,2,4-triazole (Sangon Biotech (Shanghai) Co., Ltd) medium.

**Fig. 1:**
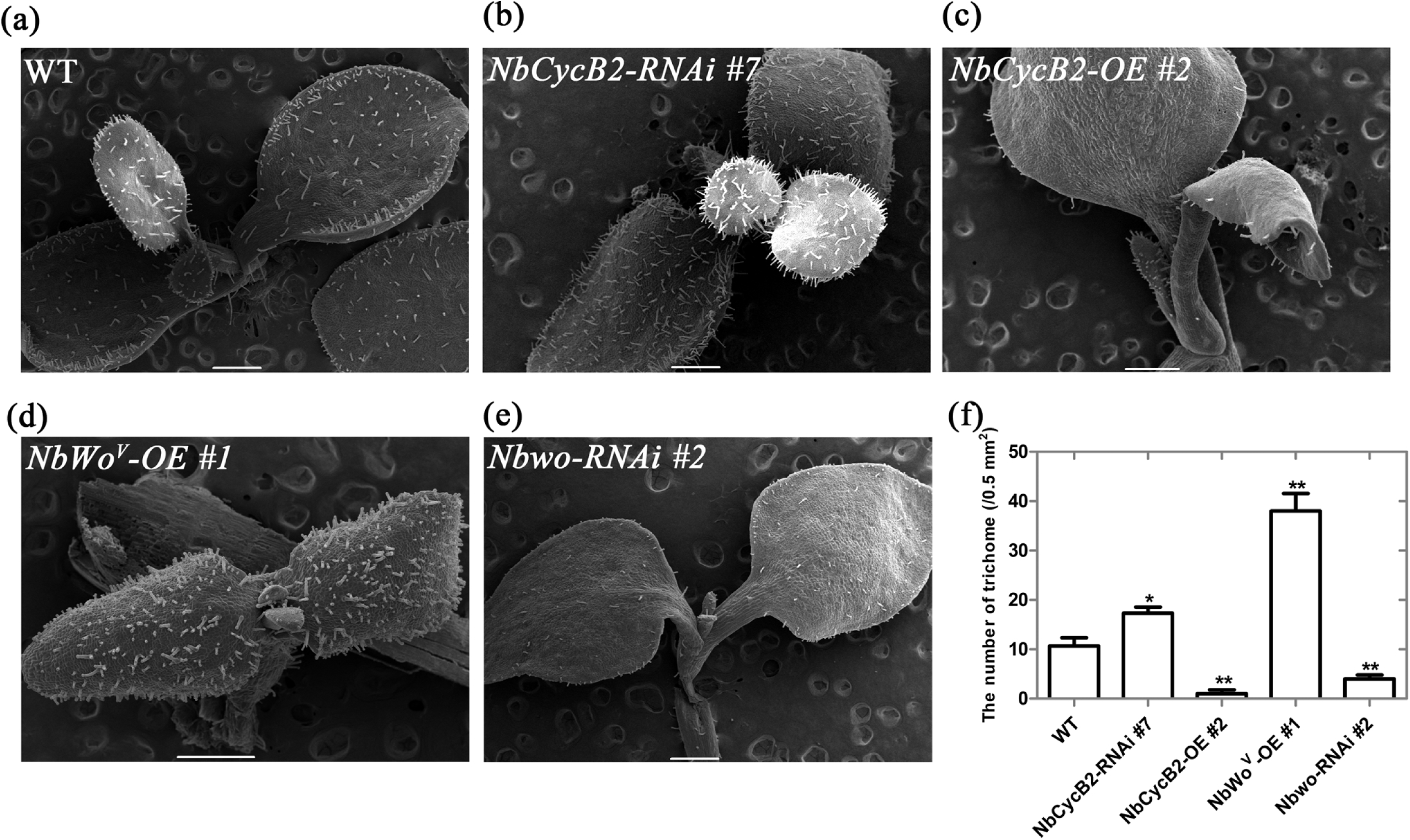
The trichome phenotypes of *NbCycB2, Nbwo* and *NbWo^v^* transgenic seedlings (a), (b), (c), (d), (e), are the trichomes SEMs of wild type, *NbCycB2-RNAi* #7 T1, *NbCycB2-OE* #2 T1, *NbWo^v^-OE* #1 T1, *Nbwo-RNAi #2* T1 10d-old-seedlings respectively. The white bar is 500 µm. (f) The trichome density of the wild type, *NbCycB2-RNAi* #7 T1, *NbCycB2-OE* #2 T1, *NbWo^v^-OE* #1 T1, *Nbwo-RNAi #2* T1 10d-old-seedlings leaves were shown. “*” indicates a difference at P<0.05 by Student’s t test compared to WT. “**” represent significant difference against WT at P < 0.01. Error bars represent SD (n = 3).

**Fig. 2:**
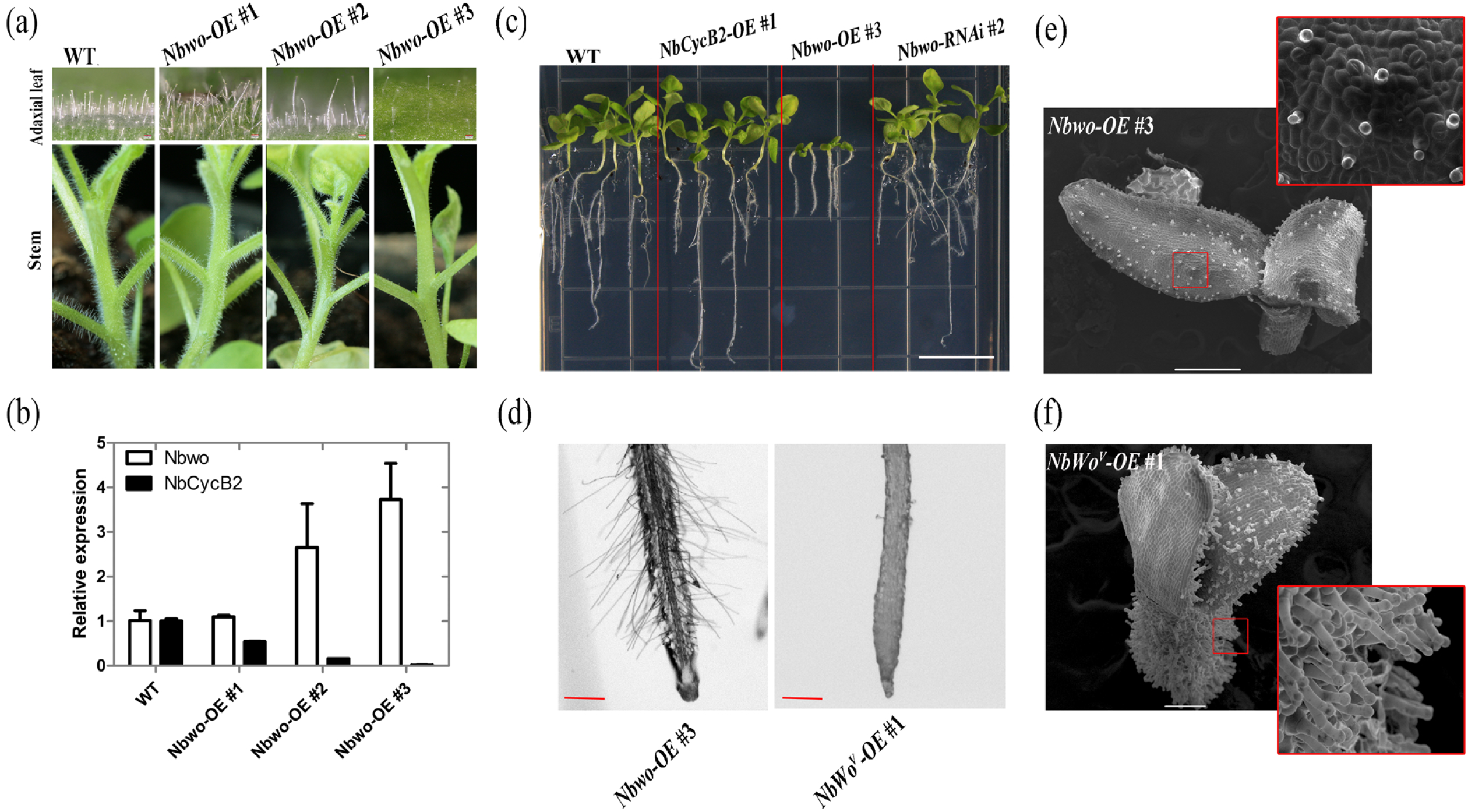
Over expression of *Nbwo* also induce the dwarf phenotype in *N. benthamiana* (a) Over expression of *Nbwo* was drivered by P35S promoter. The trichomes density reduced in the stems and leaves of transgenic lines. (b) The relative expression level of *Nbwo andNbCycB2* in the transgenic lines measured by qRT-PCR. Error bars represent SD (n = 3). (c) The root lengths were measured in the Wild type, *NbCycB2-OE* #1 T1, *Nbwo-OE* #3 T1, *Nbwo-RNAi #2* T1 two weeks-old-seedlings. The white bar is 1 cm. (d) The root hairs of *Nbwo-OE* #3 T1 and *NbWo^v^-OE* #1 T1 two weeks-old-seedlings were detected by microscope. The red bar is 1 mm. (e), (f) are the SEMs of the *Nbwo-OE* #3 T1 and *NbWo^v^-OE* #1 T1 10d-old-seedlings respectively. The white bar is 500 µm.

The yeast two-hybrid was performed to understand the interaction between *Nbwo* and *NbCycB2.* We truncated *Nbwo* into four segments containing: HD, LZ, START and SAD domains, and fused them into AD vectors to find the target region of the interaction in *Nbwo.* In addition, the CDS of *NbWo^v^* was also amplified and inserted into AD vector. Each pair of AD and BD plasmids were co-transformed into the Y2HGold yeast strain. BD-53 and AD-T constructs were co-transformed in to Y2HGold as positive control, and BD-Lam and AD-T as negative control. The transformants were then cultivated on SD/-Leu/-Trp medium (DDO) and tested on SD/-Ade/-Leu/-His/-Trp with 40 mg/L X-a-Gal (QDO/X) or SD/-Ade/-Leu/-His/-Trp with 40 mg/L X-a-Gal and 400 µg/L Aureobasidin A medium (QDO/X/A).

Yeast three-hybrid was conducted to analyze the binding competition between Nbwo LZ domain (Nbwo-LZ) and NbCycB2 to Nbwo. The Nbwo-LZ was fused with BO (Clontech). The *NbCycB2* was inserted into the downstream of methionine repressible promoter (PMet25: *NbCycB2)* in pBridge vector. The plasmid of *BD-Nbwo-LZ* was transferred with *AD-Nbwo* as positive control, and empty pBridge vector was transferred to with *AD-Nbwo* as negative control. The transformants were then tested on SD/-Ade/-Leu/-His/-Trp mediums with different concentrations of methionine (0, 250 µM).

### Bimolecular fluorescence complementation Assay (BiFc)

In order to determine the interaction of NbCycB2 and Nbwo (or NbWo^v^) in N. *benthamiana* protoplasts. The CDSs of *Nbwo, NbWo^v^* and *NbCycB2* were inserted into the pSAT6-cEYFP-C1-B vector (2×35s: YFP^c^-Nbwo, 2×35s: YFP^c^-NbWo^v^) and the pSAT6-n(1-174)EYFP-C1 vector (2×35s: YFP^n^-NbCycB2) separately (Citovsky et al., 2006). Each pair of the two plasmids were then transiently transformed into the protoplasts via PEG–calcium transfection method as described in the previous study (Yoo et al., 2007).

Moreover, to determine the interactions between Nbwo and NbCycB2 in vivo, the CDSs of *Nbwo, NbWo^v^* and *NbCycB2* were fused separately with the C-terminal fragment of YFP in p2YC vector; *Nbwo-LZ* and *NbWo^v^* were also fused with the N-terminal fragment of YFP in p2YN vector respectively (Shen et al., 2011). Different plasmid combinations were co-infiltrated into leaves of *N. benthamiana* as described in previous study (Shen et al., 2011).

The YFP fluorescence was observed by using confocal microscopy (LSM 780, Carl Zeiss, Jena, Germany). Before the observation, the transformed protoplasts were cultivation in 26 °C for 12 h, and the transformed leaves of *N. benthamiana* were cultivation in darkness for 48-72 h. Three biological repeats were observed independently for each samples.

### Dual-luciferase (Dual-Luc) assay

The regulatory effectors of 2×35s: *HA-Nbwo,* 2×35s: *HA-NbWo^v^* and 2×35s: *Flag-NbCycB2* were generated by using the DNA sequences with Ncol (5’-end) and BgIll (3’-end) cloning sites.

The firefly luciferase reporters were created by inserting the B and D fragments of the *NbCycB2* promoter, and the *Renilla* luciferase was driven by the 35S promoter in the pGreen-0800-II report vector (35S: REN-NbCycB2proB: LUC, 35S: REN-NbCycB2proD: LUC). The mutants of the *NbCycB2* promoter D fragments were also constructed using a Mutagenesis Kit (35S: REN-NbCycB2proD-m2: LUC). The regulatory effector and the reporter were used at the ratio of 5:1 or 5:5:1 for the expression test of two or three plasmids.

### lmmunoblotting, CO-IP and pull-down assay

The *N. benthamiana* leaves (∼0.5 g) were homogenized with liquid nitrogen and then solubilized with 0.4 ml of lysis buffer (25 mM Tris-HCI, 2.5 mM EDTA, pH 8.0, 0.05% v/v NP-40, 5% glycerol, 150 mM NaCl, 1 mM phenylmethylsulfonyl fluoride and 20 uM MG132) for 30 min at 4 °C. After solubilization, the protein extract was centrifuged at 13,000g for 10 min at 4 °c to separate the solubilized (supernatant) and non-solubilized material. Total protein (∼so µg) was then used for the immunoblotting assay. After SDS-PAGE separation, the proteins were electrophoretically transferred to a PVDF membrane for immunodetection.

The interaction of NbCycB2 and Nbwo dimers were determined by using co-IP assay in *N. benthamiana* protoplasts. In the expression vectors, the Nbwo protein was fused to the HA and Flag tags, respectively, and the NbCycB2 was fused to the GFP tag. Each pairs of the plasmids were transformed into the protoplasts via PEG–calcium transfection method as described in the previous study (Yoo et al., 2007). The total proteins were extracted by using lysis buffer and then incubated for 3 h at 4^°^C with 20 ul of Anti-HA Affinity Gel (Millipore). The immunoprecipitates were washed five times with lysis buffer. The isolated proteins were detected by immunoblotting with anti-Flag or anti-GFP antibodies.

In the pull-down assay, the CDSs of *Nbwo, NbWo^v^* and *NbCycB2* were respectively inserted into the pET22b and PGEX-4T-1 vectors to create the fusion proteins (His-Nbwo, His-NbWo^v^ and GST-NbCycB2), and then transformed into the *Escherichia coli* BL21 strain. The purified recombinant bait proteins (2 mg His-Nbwo or His-NbWo^v^) and 2 mg of prey proteins (GST-NbCycB2) were then mixed with 1 ml binding buffer (SO mM Tris-HCI pH 7.5, 0.6% Triton and X-100, 100 mM NaCl,). After incubation at 4 ^°^C for 2 h, 50 µl of glutathione agarose was added to the mixtures followed by the incubation for additional 1 h. The immunoprecipitates were washed five times with binding buffer. The isolated proteins were detected by immunoblotting with anti-His or anti-GST antibodies.

### Chromatin immunoprecipitation (ChIP PCR) assay

Four weeks old *NbWo^v^-OE* plants were used for ChIP assay as described in previous study (Gendrel et al., 2005). The NbWo^v^ proteins were precipitated by using HA antibody (Santa Cruz). Primers were designed to amplify 3 fragments (length ∼ 120 - 210 bp) within the 1.7k bp upstream sequence of the *NbCycB2* transcription start site (Fig. 2a), and 7 fragments within ∼ 8.7k bp of genomic DNA sequence of *Nbwo* (Fig. 7a). After the immuno-precipitation, the purified DNA was analyzed by real-time PCR with the primers of *NbCycB2* promoter and *Nbwo* genomic DNA sequence fragments (Table S1). Enrichment was calculated from the ratio of immuno-precipitated sequences.

### Phenotype observation

Images of the transgenic plants stem were obtained using Nikon camera. The observation of abaxial leaves and root hair were used an Axioplan 2 microscope (Carl Zeiss AG).

Leaf of transgenic line and wild type seedlings, which used for scanning electron microscope analysis, were fixed with 2% glutaraldehyde (0.1M phosphate buffer, PH 7.4) at 4 □ for 12h. Then, the samples were dehydrated with a series of alcohol (10%, 20%, 30%, 40%, 50%, 60%, 70%, 80%, 90% and 100%) for 20 min each time. Finally, the samples were dried in a critical point drying device (Leica EMCPD030), and coated with gold particles. The samples were observed by using a JSM-6390/LV scanning electron microscope.

### Data deposition

The sequences reported in this article have been deposited in the Sol genomic network (Fernandez-Pozo et al., 2015) (https:ljsolgenomics.net/) with the accession numbers as follows: *Nbwo* (Niben101Scf07790g01007.1), Nbwo-allele (Niben101Scf00176g11005.1), *NbCycB2* (Niben101Scf10299g00003.1), *NbCycB2-allele* (Niben101Scf10396g00002.1), *NbML1* (Niben101Scf00703g00003.1), *NbML1*-allele (Niben101Scf01158g03010.1).

## Results

### Expression and cellular analysis of *Nbwo* and *NbCycB2*

We obtained 14 Nbwo and 8 NbCycB2 protein-related sequences from online databases (Fig. S2a, b). The full-length coding sequences of *Nbwo* and *NbCycB2* obtained in the current study contained 2,199 bp and 333 bp, respectively. One allele of each of the *Nbwo* and *NbCycB2* genes were identified via BLAST with 95.82% and 95.20% identity, respectively. The amino acid analysis determined the two-point mutations at 2,084 and 2,092 of NbWo^v^ could cause two amino acid replacements in the woolly motif (Ile to Arg, Asp to Tyr, Fig. S1c). Conserved domain analysis revealed that NbCycB2 contained a WD40-like domain in the N-terminal (NbCycB2-WD40, including an EAR like motif) and a RING-like domain in the C-terminal (NbCycB2-RING) (Fig. S2d). However, no conserved domain of B-Type Cyclin protein was found in NbCycB2 protein sequences.

The visualization of subcellular localization revealed that NbCycB2, Nbwo and NbWo^v^ all localized to the nucleus (Fig. S3a). As shown in the self-activation assay, clones of AD-Nbwo and AD-NbWo^v^ can grow on QDO/X/A medium compared to the positive control, which means they have strong self-activating ability, while the mutations of woolly motif did not affect Nbwo’s transactivation ability (Fig. S3b).

As shown in Fig. S4a, the spatial expression pattern assays of *NbCycB2* and *Nbwo* indicated they were lowly expressed in roots but at a high level in the trichome containing organs. Further investigation revealed the expression of the *NbCycB2* promoter-driven GFP-GUS transgenic lines was only detected in the trichomes of leaves and stems (Fig. S4b-c). However, GUS of the Nbwo promoter-driven GUS transgenic line was strongly expressed in the basal and venous regions of young leaves (Fig. S4d).

### NbCycB2 negatively regulates trichome initiation

Most of *NbCycB2* overexpression (OE) transgenic lines underwent a dramatic reduction of trichomes on leaves and stems (Fig. S5a, Fig. lc, lf). The root length and number of branch root significantly increased in *NbCycB2-OE* lines (Fig. S6). Western blot and qRT-PCR analysis showed the *NbCycB2* transcripts significantly accumulated in *NbCycB2-OE* lines, while the expression levels of *Nbwo* and endogenous *NbCycB2* were significantly reduced (Fig. S5b, c).

In contrast, the density of trichomes increased significantly on the leaves and stems of 16 *NbCycB2* knockdown lines *(NbCycB2-RNAi)* (Fig. S5d, Fig. lb, lf). The qRT-PCR analysis showed the number of trichomes negatively correlated with the expression level of *NbCycB2* (Fig. S5e), suggesting that *NbCycB2* may play a negative role in trichome initiation.

### NbWo^v^ positively regulates trichome initiation

To confirm the function of *Nbwo,* we also generated 22 *Nbwo* knockdown transgenic plants (Nbwo-RNAi). Compared with the wild type, trichome densities were clearly decreased on the leaves and stems of most *Nbwo* knockdown plants (Fig. S7a, e). The efficiency of RNAi mediated knockdown was confirmed by qRT-PCR in two independent lines, in which the expression of *Nbwo* and *NbCycB2* was significantly reduced (Fig. S7b).

As in the previous study, dramatic increases in the density and branching of trichomes were found on the leaves and stems of *NbWo^v^-OE* plants (Fig. 1d, 1f, S7d, S7h). Our qRT-PCR assay showed the expression levels of the *NbWo^v^,* endogenous *Nbwo* and *NbCycB2* genes were all significantly upregulated in transgenic lines (Fig. S7f-g). The dwarfism phenotypes were observed in the T1 plants of *NbWo^v^-OE* (Fig. S8).

### Over-expression of *Nbwo* also induces the dwarfism

Twenty transgenic plants with overexpression of *Nbwo (Nbwo-OE)* were generated. Interestingly, the density of trichomes was shown to negatively correlate with the expression level of *Nbwo* in the T0 of *Nbwo-OE* plants (Fig. 2a, b). Additionally, the expression levels of *NbCycB2* were significantly reduced in trichome reduced plants (Fig. 2b). However, the decreased trichome phenotypes were not observed in T1 plants, but the higher expression level of exogenous *Nbwo* (for example, *Nbwo-OE #3)* also resulted in dwarfism similar to *NbWo^v^-OE* lines (Fig. 2c). Except for the dwarfism, these two lines, however had a great difference in both trichomes and root hairs. In the *Nbwo-OE #3,* the development of glandular trichomes and the development of root hairs were normal. In *NbWo^v^-OE #1,* the density of glandular trichomes as opposed to root hairs increased significantly (Fig. 2d-f).

### Nbwo and NbWo^v^ directly targeted the L1-like box of the *NbCycB2* promoter

The expression of *NbCycB2* was significantly upregulated in the *NbWo^v^-OE* lines, and decreased in the Nbwo-RNAi lines (Fig. S7b, f). These results indicated that *NbCycB2* was positively regulated by Nbwo and NbWo^v^. To test whether Nbwo directly bound to the promoter of *NbCycB2, NbWo^v^-OE (NbWo^v^* fused with HA tag) plants were analyzed by ChIP qRT-PCR assay with an HA antibody. Strong enrichment of NbWo^v^ was observed in the P2 region of the *NbCycB2* promoter in *NbWo^v^-OE* plants (Fig. 3a, b).

**Fig. 3:**
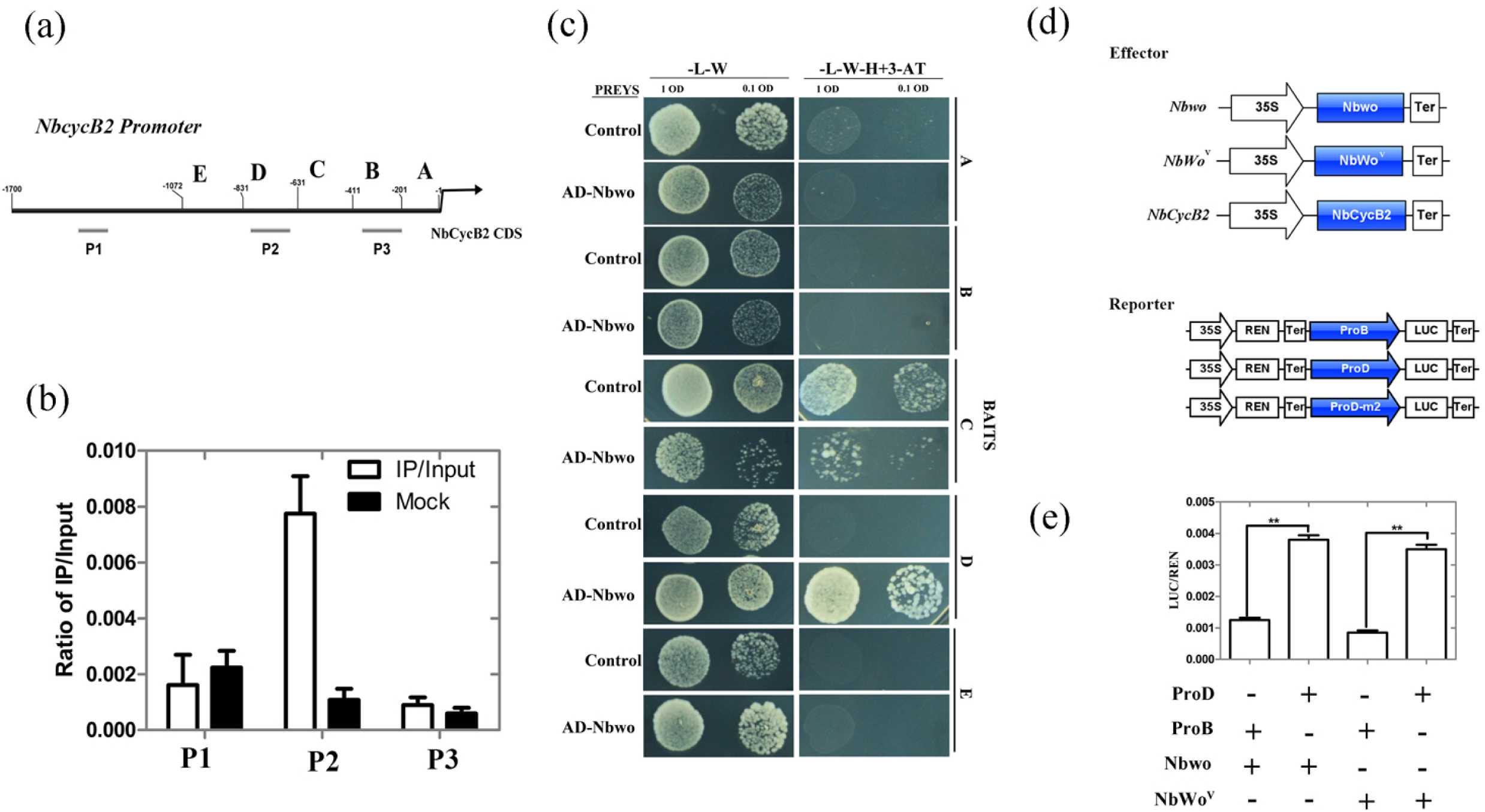
Nbwo and NbWo^v^ protein can bind *NbCycB2* promoter in vitro and vivo (a) The fragments of *NbCycB2* promoter were used to CiHP (P1, P2 and P3) and yeast one-hybrid assays (A, B, C, D and E). The numbers indicate positions of the *NbCycB2* promoter truncations. (b) Graphs show the ratio of bound promoter fragments (P1-P3) versus total input detected by qRT-PCR after immuno-precipitation in HA-NbWo^v^ by HA antibodies. Data shown are mean ± SE (n = 3). (c) One-hybrid (Y1H) assays were used to determine the interaction of *NbCycB2* promoter fragments (A, B, C, D, E) bait constructs and *AD-Nbwo* or empty pGADT7 constructs in Y187 yeast strains. (d) The schematic diagram of the effectors and reporters constructs were used in the LUC assay. (e) The relative reporter activities were measured in *N. benthamiana* protoplasts after transiently transformed the effector and reporter constructs. The relative LUC activities normalized to the REN activity are shown (LUC/REN). The difference between combinations were detected by Student’s t test. “**” present that the LUC activity is significantly different (P< 0.01). Error bars represent SD (n = 3).

To further determine the specific area of the *NbCycB2* promoter binding to Nbwo, we performed yeast one-hybrid (Y1H) assays. The five truncated fragments of *NbCycB2* promoter were shown in Fig. 3a. The yeast colonies containing NbCycB2-proD-pHIS 2 and *AD-Nbwo* constructs were grown on the selection medium with 3-aminotriazole (60 mM) (Fig. 3c).

To investigate whether Nbwo and NbWo^v^ directly affect the expression of the D fragment in vivo, we performed dual-Luc assays. The reporters 35S: REN-NbCycB2proD: LUC, 35S: REN-NbCycB2proB: LUC and the effectors were shown in Fig. 3d. As shown in Figure 3e, each pair of reporter and effector was transiently co-expressed in *N. benthamiana* protoplasts. Compared to the B fragment, when the Nbwo or NbWo^v^ was transiently co-expressed, the D fragment-driven LUC expression accumulated significantly in the protoplasts, indicating that the D fragment of the *NbCycB2* promoter should be a specific site for Nbwo and NbWo^v^binding.

Further analysis of the targeting sequence of *NbCycB2proD* revealed this sequence contained two L1-like boxes (5’-ATTTACTC-3’) (Fig. 4a). In the Y1H assay and the in vivo LUC assay, when two L1-like boxes were mutated *(NbCycB2proD-m2),* the interaction with the Nbwo protein is abolished (Fig. 4b, c). Based on these results, we inferred the L1-like boxes may be the binding target of the Nbwo and NbWo^v^ proteins.

**Fig. 4:**
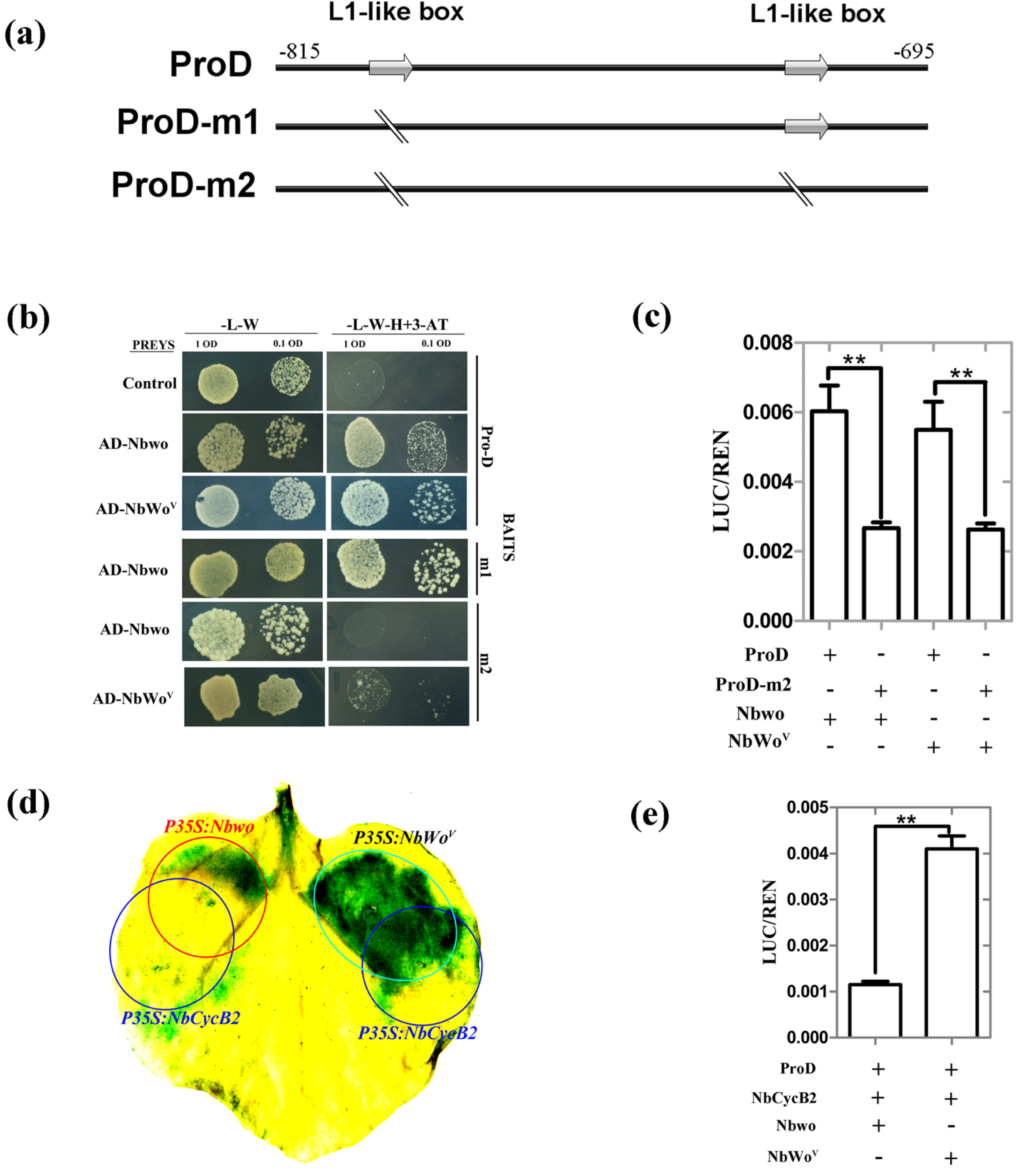
NbCycB2 can suppress the function of Nbwo, but has no effect on NbWo^v^ (a) The DNA sequences of wild type *NbCycB2* promoter (ProD) and mutant fragments were shown. L1-like boxes were indicated by using gray arrow. The mutated bases in the m1 and m2 sequences are shown by a dividing line. (b) One-hybrid (Y1H) assays were used to detected the interaction between mutant D fragments of *NbCycB2* promoter bait constructs and *AD-Nbwo* or *AD-NbWo^v^* in Y187 yeast strains, the empty pGADT7 construct as the control. (c) Relative reporter activity was measured in *N. benthamiana* protoplasts after transiently co-transformed the effector and reporter constructs. The relative LUC activities normalized to the REN activity are shown (LUC/REN). “**” present that the LUC activity has significant difference between the D and m2 fragments reporter when their co-transformed with *Nbwo* or *NbWo^v^* effector respectively (P< 0.01, Student’s t test). Error bars represent SD (n = 3). (d) The GUS staining of the *proNbCycB2: GFP-GUS* transgenic line leaves, when co-expressing *P35S: Nbwo* and *P35S: NbCycB2, P35S: NbWo^v^* and *P35S: NbCycB2.* The area of red circle was injected with *P35S: Nbwo* GV3101 strain; the area of Green circle was injected with *P35S: NbWo^v^* GV3101 strain; the area of blue circles were injected with *P35S: NbCycB2* GV3101 strain. (e) The LUC activity of co-transforming the effector and reporter constructs were measured. “**” present that the LUC activity is significantly different between *Nbwo* and *NbWo^v^* effector when they co-transformed with *NbCycB2* effector respectively (P< 0.01, Student’s t test). Error bars represent SD (n = 3).

### NbCycB2 represses the activity of Nbwo rather than NbWo^v^

Since the *NbCycB2-OE* and *Nbwo-RNAi* transgenic lines shared the non-trichome phenotype, and the expression of *Nbwo* was not enhanced in the *NbCycB2-RNAi* lines (Fig. S5e), we suspect that NbCycB2 could affect Nbwo transactivation ability at protein levels. To confirm this, constructs of overexpression *Nbwo, NbWo^v^* and *NbCycB2* were transiently co-expressed in the leaves of the *NbCycB2pro: GFP-GUS* transgenic line using the *Agrobacterium-mediated* method (Fig. 4d). The GUS induced by the transient expression of *Nbwo-OE* was only detected in the area where there was no *NbCycB2-OE* Expression. However, the transient expression of *NbWo^v^* caused a strong GUS staining which is independent of *NbCycB2-OE* expression. These results were further supported by the LUC assay, in which co-expression with *NbCycB2* could clearly repress the transactivation activity of Nbwo, but did not affect the NbWo^v^ protein (Fig. 4e). These results suggest that NbCycB2 may act as a negative regulator of Nbwo rather than NbWo^v^.

### NbCycB2 suppresses the transactivation ability of Nbwo via the direct interaction

The interaction between NbCycB2 and Nbwo was reported in a previous study (Yang et al., 2011). To explore which domain was involved in the physical interaction between NbCycB2 and Nbwo, four truncated fragments of Nbwo containing HD, LZ, START and SAD were used (Fig. 5a). Y2H assays suggested that the LZ domain of Nbwo interacted with NbCycB2 (Fig. 5b). The BiFC assay was further used to verify the interaction between the Nbwo LZ domain and NbCycB2 in vivo (Fig. 5c).

**Fig. 5:**
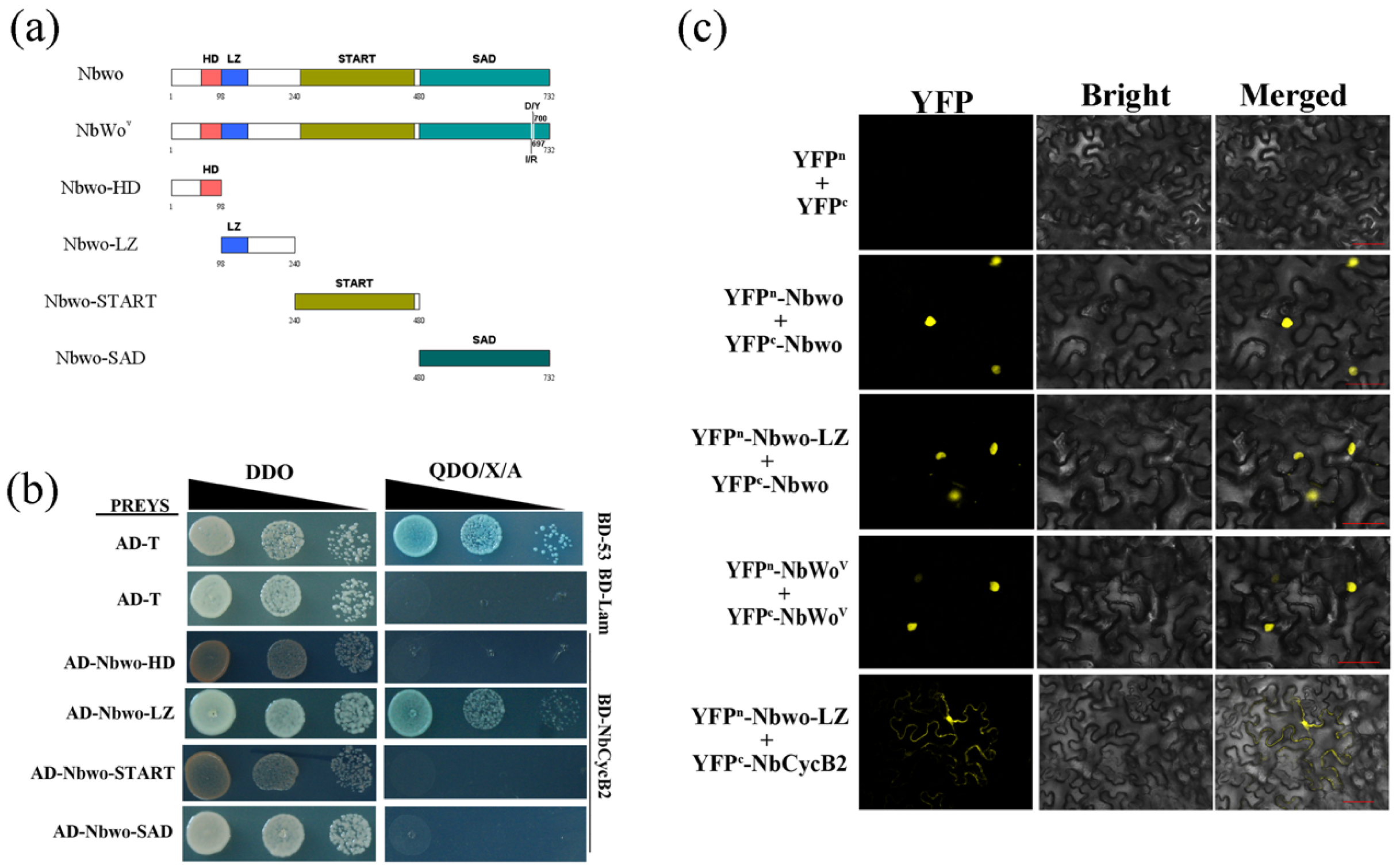
NbCycB2 protein interact with LZ domain of the Nbwo protein in vitro and vivo (a) Schematic diagrams of Nbwo protein domain constructs. The numbers indicate positions of the first and the last amino acid of the Nbwo truncations. (b) Interaction between NbCycB2 and domains of Nbwo protein were determined by YH2 system. (c) The interaction between Nbwo and NbCycB2 or itself were demonstrated by BiFC assay. Each indicated pair of constructs were co-infiltrated into leaves of *N. benthamiana* (Bar, 50 µm).

To further examine if NbCycB2 also interacts with NbWo^v^, we used yeast two-hybrid (Y2H) assays. Our results indicate that NbCycB2 physically interacts with Nbwo but did not interact with NbWo^v^, suggesting that the interaction between NbCycB2 and Nbwo can be reduced by mutations in the woolly motif (Fig. 6a). Consistent with the Y2H result, the interaction between NbCycB2 and Nbwo or NbWo^v^ was further confirmed by the pull-down assay (Fig. 6b) and the bimolecular fluorescence complementation (BiFC) assay (Fig. S10a).

**Fig. 6:**
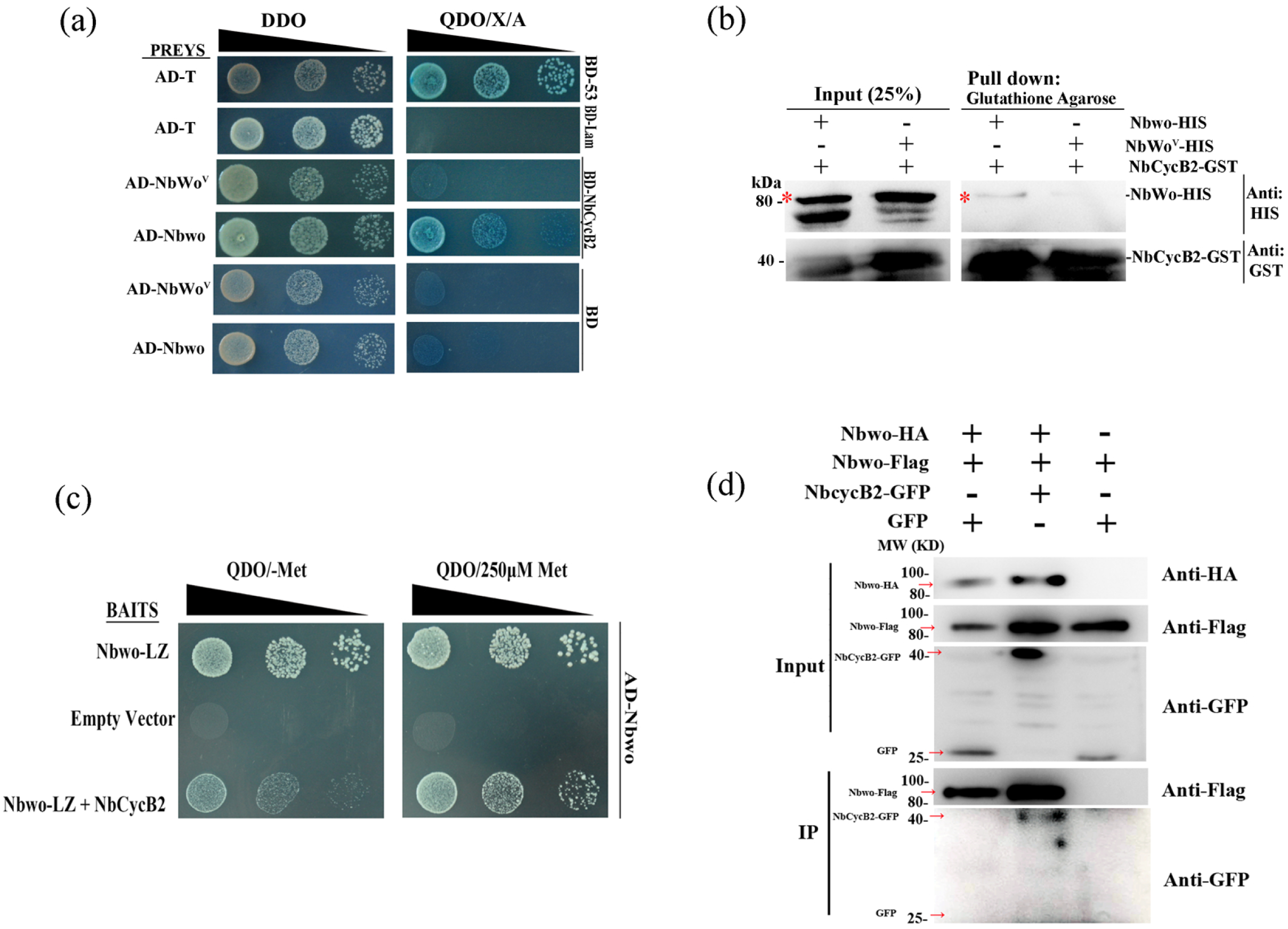
Mutation of woolly motif reduce the interaction between Nbwo and NbCycB2 (a) Interaction between NbCycB2 and Nbwo or NbWo^v^ proteins were determined by YH2 system. Blue clones grown on the QDO/X/A medium indicates positive protein-protein interactions. (b) The pull down assay between Nbwo or NbWo^v^ with NbCycB2 proteins. Only recombinant HIS-Nbwo protein can co-precipitate with GST-NbCycB2 protein. (c) The competitive binding between NbCycB2 and LZ domain, LZ domain and Nbwo were determined by Yeast three-hybrid assays. (d) The co-IP assay between NbCycB2 and Nbwo dimers. The total protein immunoprecipitation was performed by using anti-HA beads.

### Nbwo was restrained by NbCycB2 through forming a homodimer

It is known that the HD-Zip proteins bind to DNA as dimers by the LZ domain (Ariel et al., 2007). To verify whether Nbwo can be dimerized via the LZ domain, we first demonstrated in the Y2H that the LZ domain could bind to the Nbwo protein (Fig. S10b). In addition, BiFC assays were used to confirm the interaction between the LZ domain and Nbwo (or NbWo^v^) proteins in vivo (Fig. 5c).

To test whether the NbCycB2 could bind at the Nbwo homodimer, we conducted the yeast three-hybrid (Y3H) and co-IP assays. As shown in Fig. 6c-d, NbCycB2 could interact with the Nbwo dimers.

### Nbwo could bind to its own genomic DNA

The endogenous expression level of *Nbwo* reduced in *NbCycB2-OE* lines (Fig. S5b) and increased in *NbWo^v^-OE* plants (Fig. 1d), indicated that Nbwo may be able to regulate self expression. In order to prove this hypothesis, ChIP assay was carried out to check whether Nbwo could bind to its genomic DNA sequence in the leaf of *NbWo^v^-OE* transgenic line. Interestingly, enrichment of NbWo^v^ was found in the T5 fragment in *NbWo^v^-OE* plants (Fig. 7a, b). This result was further demonstrated by using Y1H assay. Only the clones with AD-Nbwo (or AD-NbWo^v^) and Nbwo-G3-pHIS 2 constructs could grow on the resistant medium, suggesting that Nbwo or NbWo^v^ could bind to the G3 fragments (include T5 fragment, Fig. 7a) of its own genomic DNA sequences (Fig. 7c).

**Fig. 7:**
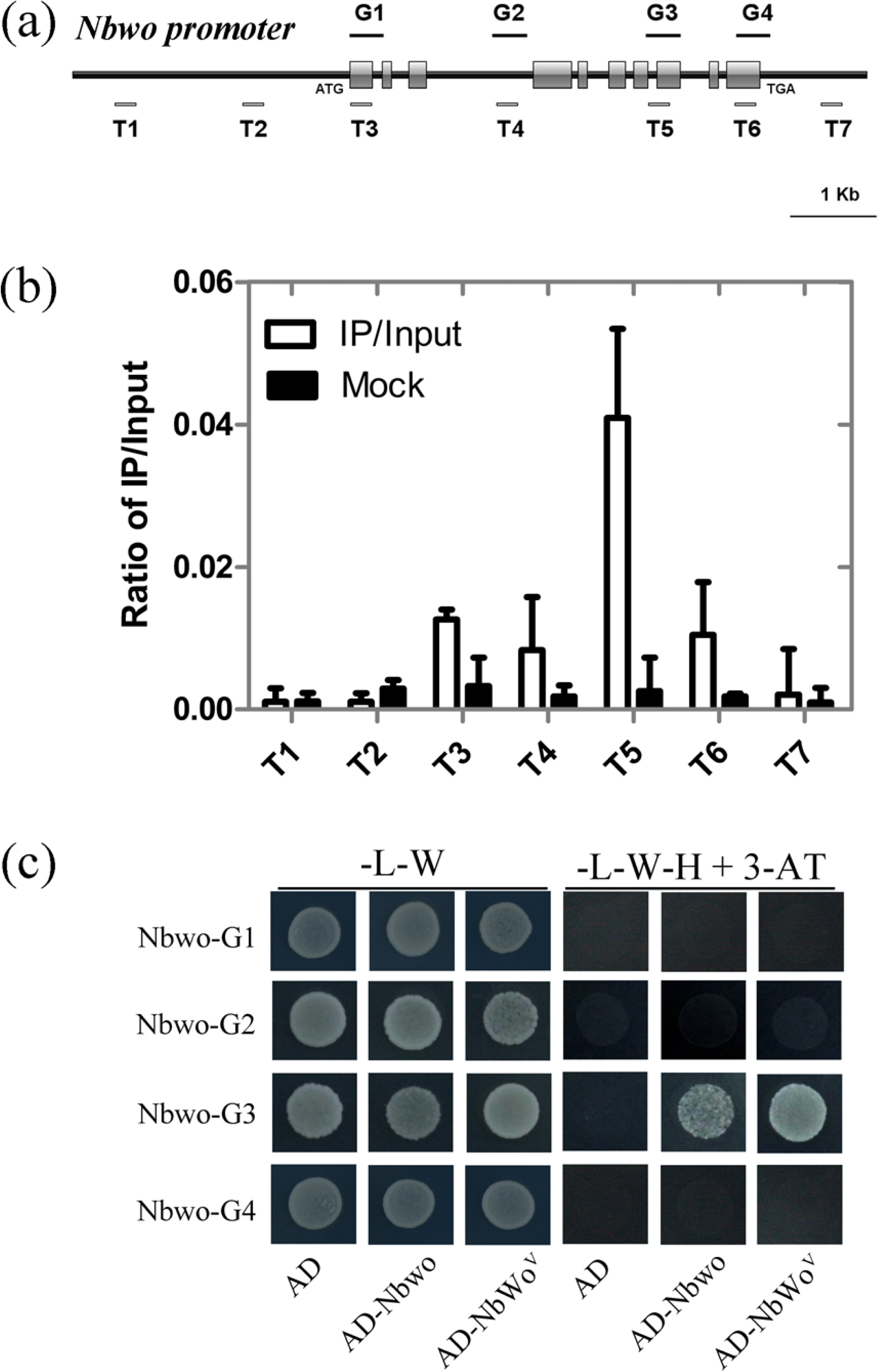
Nbwo and NbWo^v^ can bind at itself genomic DNA sequences (a) The fragments of *Nbwo* genomic sequences were used to CHiP (T1, T2, T3, T4, TS, T6 and T7) and yeast one-hybrid assays (G1, G2, G3 and G4). The bar present 1 kb DNA sequences. (b) Graphs show the ratio of bound genomic fragments (T1-T7) versus total input detected by real-time PCR after immuno-precipitation in *NbWo^v^-OE* lines by HA antibodies. Data shown are mean ± SE (n = 3). (c) One-hybrid (Y1H) assays were used to determine the interaction of *Nbwo* genomic sequence fragments (G1, G2, G3, G4) bait constructs and AD-Nbwo, AD-NbWo^v^ or empty pGADT7 constructs in Y187 yeast strains. The clones grown on the SD/-Leu/-His/-Trp (-L-W-H) with 60 mM 3-AT medium indicates the interaction between DNA fragment and Nbwo or NbWo^v^ proteins.

### Overexpression of *NbCycB2* can reduce the dwarf phenotype of *Nbwo-OE* plants

To determine whether *NbCycB2* can inhibit the transactivation activity of *Nbwo* in vivo, we crossed *NbCycB2-OE #2* T1 plant to *Nbwo-OE #3* T0 plants. As shown in Fig. 8 a-b, the dwarf and short-root phenotypes of *Nbwo-OE #3* were indeed reduced by the *NbCycB2-OE #2.* The crossed F1 plants were tested using PCR (Fig. 8 c), and the expression of *NbCycB2* and *Nbwo* was also verified by qRT-PCR. Compared with T1 *Nbwo-OE* #3 plants, the balance between *NbCycB2* and *Nbwo* expression was restored in *NbCycB2-OE* #2 and *Nbwo-OE* #3 crossed F1 plants (Fig. 8 d).

**Fig. 8:**
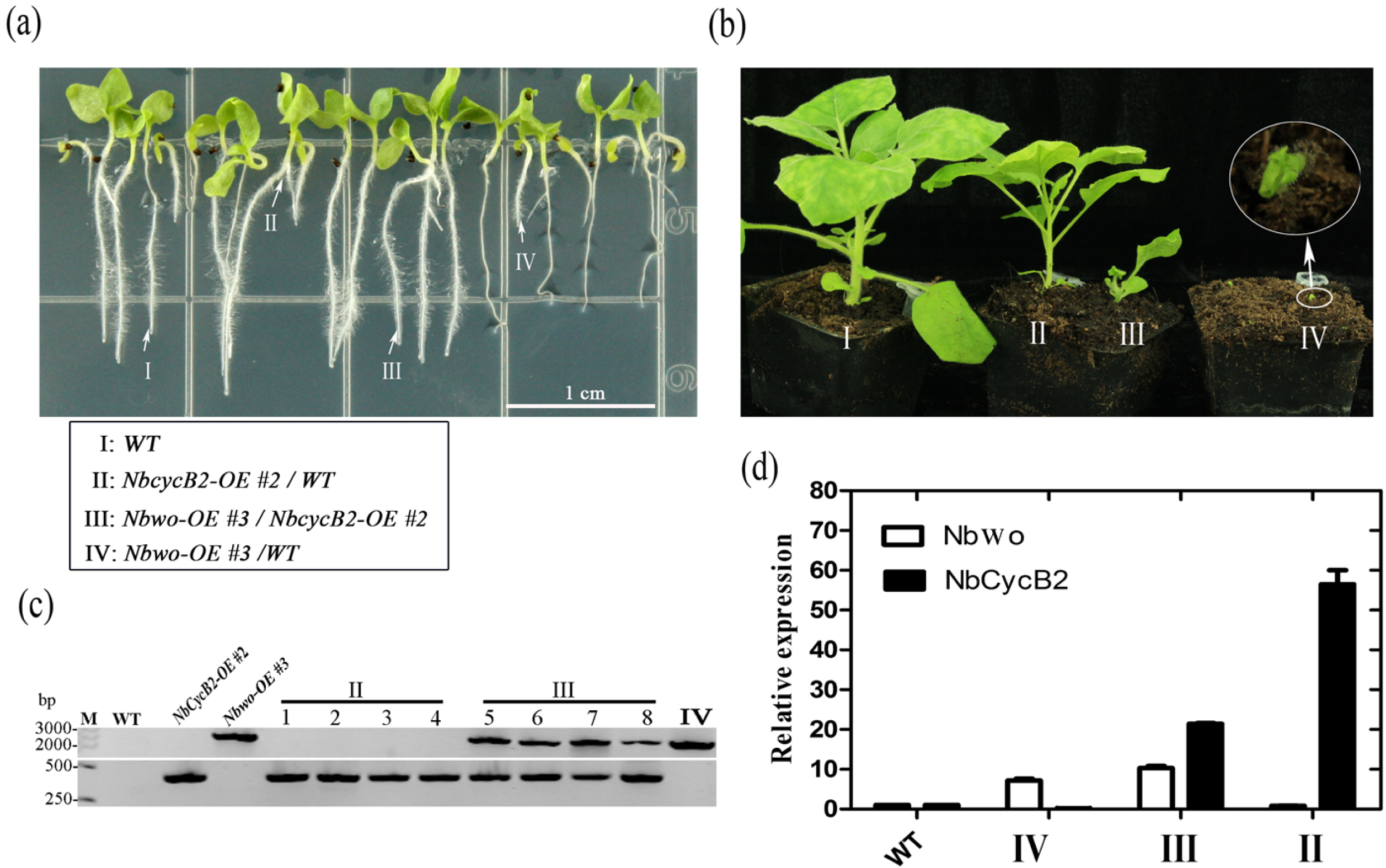
Hybridization between *Nbwo-OE* and *NbCycB2-OE* plants (a), (b) The phenotype of *NbCycB2-OE* #2 and *Nbwo-OE* #3 hybridization F1 two weeks old (a) and mature (b) plants were shown. “I” present wild type *N. benthamiana,* “II” present *NbCycB2-OE/WT* hybrid T1 plants, “III” present *NbCycB2-OE #2* and *Nbwo-OE #3* hybridization F1 plants, “IV” present *Nbwo-OE #3/WT* hybrid T1 plants. (c) F1 plants were tested by using PCR. Wild type as a negative control, *Nbwo-OE* #3 and *NbCycB2-OE* #2 as positive controls. No DNA band was detected in the wild type *N. benthamiana.* A ∼330 bp DNA band was detected in *NbCycB2-OE* #2 lines. A ∼2199 bp DNA band was detected in *Nbwo-OE #3* lines. In contrary, two bands were detected in *NbCycB2-OE #2* and *Nbwo-OE #3* hybridization F1 plants. (d) The relative expression levels of *Nbwo* and *NbCycb2* were measured by qRT-PCR in F1 plants. Data are given as means SD (n = 3).

## Discussion

### *NbCycB2* negatively regulates trichome initiation

It is well known that ectopic expression of a constitutive active B-type cyclin promoted the unicellular trichome differentiate to multicellular in *Arabidopsis* (Schnittger et al., 2002, 2005). The *SlCycB2* gene has been reported to be a hypothesis B-type cyclin gene participating in trichome formation in tomato (Yang et al., 2011; Gao et al., 2017). However, the function of *SlCycB2* during trichome development has not been well-studied.

In this study, we found that overexpression of *NbCycB2* caused a non-trichome phenotype, whereas inhibition of *NbCycB2* significantly increased the density of trichomes instead of branching on stems and leaves (Fig.1b, S5). Consistent with the qRT-PCR results (Fig. S4a), GUS staining assay indicated *NbCycB2* is specifically expressed in the trichomes of leaves and stems (Fig. S4b-c), and *Nbwo* is expressed in the basal and venous regions of young leaves (Fig. S4d). These results suggested NbCycB2 serves as a negative regulator of trichome initiation. None of B-type cyclin conserve domains were found in the SlCycB2 and NbCycB2 protein sequences (Fig. S2d). Thus, whether SlCycB2 has the function of B type cyclin protein requires further study.

### NbWo^v^ and Nbwo directly regulate the expression of *NbCycB2* through binding to the Ll-like boxes in the promoter

In previous study, *SlCycB2* has been reported to be indirectly regulated by *Wo^v^* (Yang et al., 2011; Yang et al., 2015). The expression level of *NbCycB2* was upregulated in overexpression of *NbWo^v^* plants and downregulated in *Nbwo-RNAi* lines (Fig. S7), indicating that *NbCycB2* may be the downstream gene of *Nbwo.* Additionally, the D fragments of the *NbCycB2* promoter have been shown to be the binding target of Nbwo and NbWo^v^ through ChIP, Y1H and LUC assays (Fig. 3). Mutation of the two L1-like box sequences in *NbCycB2proD* inhibited the binding of Nbwo or NbWo^v^ in vitro and vivo (Fig. 4b). Therefore, we proved that *NbCycB2* was directly regulated by Nbwo or NbWo^v^ through the binding of L1-like boxes in the *NbCycB2* promoter (Fig. 4a-c). In addition, we also proved that Nbwo and NbWo^v^ can self-regulate the endogenous expressions by binding to its own genomic DNA sequence by ChIp and YIH assay (Fig. 7).

### The increase of trichome density and plant dwarfism was regulated by Nbwo through different pathways

The *Wo* homozygous plants have been shown to cause the embryo lethality in seeds (Yang et al., 2011). We also found the overexpression of *Nbwo* or *NbWo^v^* genes causes abnormal embryonic development and results in dwarf phenotype in its offspring (Fig. 2c, S6). However, the trichome phenotype of *Nbwo-OE* and *NbWo^v^-OE* was completely different (Fig. 2e and 2f), suggesting that Nbwo was involved in both the regulation of the development of trihcome and embryo. The trichomes’ density was decreased with the expression of Nbwo in T0 of *Nbwo-OE* lines (Fig.2a and 2e), which suggested that high expression of *Nbwo* in wild type background could repress trichome development. Thus, we suspected the woolly motif represses *Nbwo* activity. However, the trichome density was not increased when the SAD domain deleted *Nbwo* CDS (including woolly motif) was overexpressed in the wild type plants (Fig. S9), which suggesting that woolly motif did not have repressing activity. Therefore, the decreased trichome density in high expression of *wo* gene plants requires further investigation.

### NbCycB2 represses the transactivation activity of Nbwo at protein level

The *NbCycB2* gene was proven to be directly regulated by Nbwo in our study. However, the similar non-trichome phenotypes of *NbCycB2-OE* and *Nbwo-RNAi* transgenic lines were found (Fig. S5a, S7a). Additionally, the expression of *Nbwo* was not increased in *NbCycB2-RNAi* plants (Fig. S5e). These results suggested that NbCycB2 might repress the transactivation activity of Nbwo at protein levels. The GUS activity of the *NbCycB2pro: GFP-GUS* transgenic line was upregulated by the expression of *Nbwo* and inhibited by the co-expression of *NbCycB2* (Fig. 4d), and the same result was found in the LUC assay (Fig. 4e). Additionally, Hybridization with *NbCycB2-OE* can attenuate the dwarf phenotype of T1 *Nbwo-OE* (Fig. 8a-b). Further studies revealed that the expression of endogenous *NbCycB2* and *Nbwo* were reduced in the *NbCycB2-OE* lines (Fig. S5b). They were shown to be downstream regulatory genes of Nbwo (Figures 3 and 7). These results altogether supported our hypotheses that NbCycB2 may act as a negative regulator of Nbwo at protein levels.

In a previous study, SlCycB2 was reported to interact with wo protein (Yang et al., 2011). Further investigation of the interaction between Nbwo and NbCycB2 revealed the dimerized LZ domain of Nbwo binds to NbCycB2 (Fig. 5b, c). Through Y2H, BiFC, Y3H and co-IP assays, we also found that Nbwo could form a homodimer through the LZ domain, and the NbCycB2 protein could bind to LZ domain of Nbwo dimers (Fig. 5b-c, 6 c-d, S10b). These results indicated that NbCycB2 may bind to Nbwo protein via its LZ domain to form a complex, which inhibits its transactivation ability. However, further study is required to determine whether *NbCycB2* functions similarly to its homologue gene --AT5G06270.1 which also interacts with the co-repressor TOPLESS (Long et al., 2006; Szemenyei et al., 2008; Pauwels et al., 2010; Wu and Citovsky, 2017a).

### The interaction between Nbwo and NbCycB2 was blocked by the mutation in the Nbwo woolly motif

In *Arabidopsis thaliana,* the feedback loop regulation mechanisms of the R3 MYB (TRY, CPC and so on) is through competitively binding to GL3/EGL3 to form a non-functional trimeric protein complex (MYB-bHLH-WDR), inhibiting the formation of trichomes (Wang et al., 2008; Wester et al., 2009). Feedback loop regulation has been reported as an effective strategy to maintain normal organism development by many HD-ZIP proteins (Ohgishi et al., 2001; Williams and Fletcher, 2005; Kim et al., 2008; San-Bento et al., 2014). However, the trichome formation was not repressed by the high expression level of *NbCycB2* in the *NbWo^v^-OE* plants (Fig.1d, S7d, f), which suggest that the negative effect of *NbCycB2* could be eliminated by the mutation in *NbWo^v^.* The LUC assay and GUS activity assay in the leaves of *NbCycB2pro: GFP-GUS* transgenic lines also supported this conclusion (Fig. 4d, e).

Further investigation demonstrated the interaction between NbCycB2 and Nbwo could be blocked by the mutation of woolly motif in NbWo^v^ protein in vitro and in vivo (Fig. 6a-b, S10a), indicating that NbWo^v^ abolishes interaction with NbCycB2 to prevent inhibition of NbCycB2. The high expressions of *NbCycB2* and endogenous *Nbwo* in *NbWo^v^-OE* lines also supported this conclusion (Fig. S7f).

In summary, through this study, we found NbCycB2 is specifically expressed in the trichomes of *N. benthamiana* and negatively affects trichome formation. Further study revealed the Nbwo and NbWo^v^ were demonstrated to directly regulate *NbCycB2* and *Nbwo* expressions by binding to the L1-like box in the *NbCycB2* promoter and its own genomic DNA sequences. In addition, the NbCycB2 protein may via binding to the LZ domain of Nbwo dimers, which represses the activity of Nbwo and reduces the expression of *Nbwo* downstream genes, eventually leading to the inhibition of trichome initiation. By contrast, the interaction between NbCycB2 and Nbwo could be blocked by the mutation in woolly motif (NbWo^v^), which prevents the repression by NbCycB2, and results in the dramatic increase of trichome density and branching. In previous studies, since *SlCyc82* is highly expressed in the *Wo* (wo gain-of-function mutant alleles) and *Wo^v^* lines and underexpressed in the *Wo-RNAi* lines, SlCycB2 is believed to promote the development of type I trichomes in tomato (Yang et al., 2011). However, the function and detailed molecular mechanisms of NbCycB2 in regulating trichome development have not been studied. Our findings provide further insights into the regulatory network of *NbCycB2* and *Nbwo* or *NbWo^v^* in multicellular trichomes development of *N. benthamiana* (Fig. S11).

## Acknowledgements

We would like to thank Prof. Xi Huang for kindly providing pBridge and pGreen-0800-II report vectors and thank Dr. Yongjia Zhong, Fujian Agriculture and Forestry university, for kindly providing p2YN and p2YC vectors, and thank Prof. Tao Huang, Prof. Yi Tao and Prof. Hongrui Wang for the help in the implement of experiments. The authors declare no competing financial interests. This study was supported by the State Tobacco Monopoly Administration of China [grant No. 110201401003 (JY-03)], XMU Training Program of Innovation and Enterpreneurship for Undergraduates (2016Y0635) and Guizhou Science and Technology Major Project [Grant No: (2019)3001-2].

## Author contributions

Liang Chen, Hong Cui, Shuang Wu and Minliang Wu conceived and designed the experiments. Minliang Wu, Yuchao Cui, Li Ge, Lipeng Cui, Zhichao Xu, Hongying Zhang, Zhaojun Wang and Dan Zhou performed the experiments and analysed the results. Minliang Wu and Shuang Wu wrote the main manuscript text. Liang Chen supervised the project.

## Supplemental Data

Supplemental Fig. S1: The Scanning electron micrographs (SEMs) of trichomes in the leaf of *N. benthamiana*.

Supplemental Fig. S2: Sequence analysis of *Nbwo, NbCycB2* and their similar proteins.

Supplemental Fig. S3: The subcellular localization and auto activation test of *NbCycB2, Nbwo* and *NbWo^v^*.

Supplemental Fig. S4: The expression pattern of *NbCycB2* and *Nbwo* in *N. benthamiana*.

Supplemental Fig. S5: Overexpression of *NbCycB2* and RNA interference of *NbCycB2* in *N. benthamiana*.

Supplemental Fig. S6: The root phenotypes of wild type, *NbCycB2-RNAi* #7 T1, *NbCycB2-OE* #2 T1, *NbWo^v^-OE* #1 T1, *Nbwo-RNAi#2* T1 seedlings

Supplemental Fig. S7: RNA interference of *Nbwo* and overexpression of *NbWo^v^* in *N. benthamiana*.

Supplemental Fig. S8: The phenotype of *NbWo^v^-OE* lines.

Supplemental Fig. S9: The phenotype of overexpressing *Nbwo-SAD-mutant* in *N. benthamiana*.

Supplemental Fig. S10: The interaction between Nbwo and NbCycb2, Nbwo and Nbwo LZ domain.

Supplemental Fig. S11: A simplified model for regulation between *Nbwo* and *NbCycb2*.

Supplemental Table S1: Primers used in this study.

## Reference

Abe, M., Takahashi, T., and Komeda, Y. (2001). Identification of a cis-regulatory element for L 1 layer-specific gene expression, which is targeted by an L1-specific homeodomain protein. The Plant Journal 26, 487–494.

Ariel, F.D., Manavella, P.A., Dezar, C.A., and Chan, RL. (2007). The true story of the HD-Zip family. Trends in plant science 12, 419–426.

Bombarefy, A, Rosli, H.G., Vrebalov, J., Moffett, P., Mueller, L.A, and Martin, G.B. (2012). A draft genome sequence of Nicotiana benthamiana to enhance molecular plant-microbe biology research. Molecular Plant-Microbe Interactions 25, 1523–1530.

Chen, S., Songkumam, P., Liu, J., and Wang, G.L. (2009). A versatile zero background T-vector system for gene cloning and functional genomics. Plant physiology 150, 1111–1121.

Citovsky, V., Lee, L-Y., Vyas, S., Glick, E., Chen, M.-H., Vainstein, A, Gafni, Y., Gefvin, S.B., and Tzfira, T. (2006). Subcellular localization of interacting proteins by bimolecular fluorescence complementation in planta. Journal of molecular biology 362, 1120–1131.

Crooks, G.E., Hon, G., Olandonia, J.M., and Brenner, S.E. (2004). Weblogo: a sequence logo generator. Genome research 14, 1188–1190.

Cui, Y., Rao, S., Chang, B., Wang, X., Zhang, K, Hou, X., Zhu, X., Wu, H., Tian, Z, Zhao, Z., Yang, C., and Huang, T. (2015). Atla 1 protein initiates IRES-dependent translation of WUSCHEL mRNA and regulates the stem cell homeostasis ofArabidopsisin response to environmental hazards. Plant, Cell & Environment 38, 2098–2114.

Fernandez-Pozo, N., Menda, N., Edwards, J.D., Saha, S., Teele, I.Y., Strickler, S.R, Bombarely, A, Fisher-York, T., Pujar, A, Foerster, H., Yan, A., and Mueller, L.A. (2015). The Sol Genomics Network (SGN)-from genotype to phenotype to breeding. Nucleic acids research 43, D1036–D1041.

Freeman, B.C., and Beattie, G.A (2008). An overview of plant defenses against pathogens and herbivores. The Plant Health Instructor http://dx.doi.org/10.1094/PHI-I-2008-0226-01.

Gan, L., Xia, K, Chen, J.G., and Wang, S. (2011). Functional characterization of TRICHOMELESS2, a new single-repeat R3 MYB transcription factor in the regulation of trichome patterning in Arabidopsis. BMC plant biology 11, 1–12.

Gao, S., Gao, Y., Xiong, C., Yu, G., Chang, J., Yang, Q., Yang, C., and Ye, Z. (2017). The tomato B-type cyclin gene, SlCycB2, plays key roles in reproductive organ development, trichome initiation, terpenoids biosynthesis and Prodenia litura defense. Plant Science 262, 103–114.

Gendrel, AV., Lippman, Z., Martienssen, R., and Colot, V. (2005). Profiling histone modification patterns in plants using genomic tiling microarrays. Nature Methods 2, 213–218.

Glas, J.J., Schimmel, B.C., Alba, J.M., Escobar-Bravo, R., Schuurink, R.C., and Kant, M.R. (2012). Plant glandular trichomes as targets for breeding or engineering of resistance to herbivores. International journal of molecular sciences 13, 17077–17103.

Goodin, M.M., Zaitlin, D., Naidu, RA, and Lommel, SA (2008). Nicotiana benthamiana: its history and future as a model for plant-pathogen interactions. Molecular plant-microbe interactions 21, 1015–1026.

Grebe, M. (2012). The patterning of epidermal hairs in Arabidopsis--updated. Current opinion in plant biology 15, 31–37.

Guo, C., Luo, C., Guo, L., Li, M., Guo, X, Zhang, Y., Wang, L., and Chen, L. (2016). OsSIDP366, a DUF1644 gene, positively regulates responses to drought and salt stresses in rice. Journal of integrative plant biology 58, 492–502.

Hollósy, F. (2002). Effects of ultraviolet radiation on plant cells. Micron 33, 179–197.

Huchelmann, A, Boutry, M., and Hachez, C. (2017). Plant Glandular Trichomes: Natural Cell Factories of High Biotechnological Interest. Plant physiology 175, 6–22.

Jefferson, R.A, Kavanagh, T.A, and Bevan, M.W. (1987). GUS fusions: beta-glucuronidase as a sensitive and versatile gene fusion marker in higher plants. The EMBO journal 6, 3901.

Kang, J.H., Campos, M.L., Zemelis-Durfee, S., Al-Haddad, J.M., Jones, AD., Telewski, F.W., Brandizzi, F., and Howe, G.A (2016). Molecular cloning of the tomato Hairless gene implicates actin dynamics in trichome-mediated defense and mechanical properties of stem tissue. Journal of experimental botany 67, 5313–5324.

Kim, Y.S., Kim, S.G., Lee, M., Lee, I., Park, H.Y., Seo, P.J., Jung, J.H., Kwon, E.J., Suh, S.W., Paek, KH., and Park, C.M. (2008). HD-ZIP III activity is modulated by competitive inhibrtors via a feedback loop in Arabidopsis shoot apical meristem development. The Plant cell 20, 920–933.

Kirik, V., Simon, M., Huelskamp, M., and Schiefelbein, J. (2004). The ENHANCER OF TRY AND CPC1 gene acts redundantly with TRIPTYCHON and CAPRICE in trichome and root hair cell patterning in Arabidopsis. Developmental Biology 268, 506–513.

Letunic, I., and Bork, P. (2018). 20 years of the SMART protein domain annotation resource. Nucleic acids research 46, D493–D496.

Long, J.A, Ohno, C., Smith, Z.R, and Meyerowitz, E.M. (2006). TOPLESS Regulates Apical Embryonic Fate in Arabidopsis 312, 1520–1523.

Mauricio, R, and Rausher, M.D. (1997). Experimental manipulation of putative selective agents provides evidence for the role of natural enemies in the evolution of plant defense. Evolution, 1435–1444.

Ogawa, E., Yamada, Y., Sezaki, N., Kosaka, S., Kondo, H., Kamata, N., Abe, M., Komeda, Y., and Takahashi, T. (2015). ATML1 and PDF2 Play a Redundant and Essential Role in Arabidopsis Embryo Development. Plant & cell physiology 56, 1183–1192.

Ohgishi, M., Oka, A, Morelli, G., Ruberti, I., and Aoyama, T. (2001). Negative autoregulation of the Arabidopsis homeobox gene ATHB-2. The Plant Journal 25, 389–398.

Oppenheimer, D.G., Hennan, P.L., Shan, S., Esch, J., and Marks, M.D. (1991). A myb gene required for leaftrichome differentiation in Arabidopsis is expressed in stipules. Cell 67, 483–493.

Pattanaik, S., Patra, B., Singh, S.K, and Yuan, L. (2014). An overview of the gene regulatory network controlling trichome development in the model plant, Arabidopsis. Frontiers in Plant Science 5, 259.

Pauwels, L., Barbero, G.F., Geerinck, J., Tilleman, S., Grunewald, W., Perez, AC., Chico, J.M., Bossche, RV., Sewell, J., Gil, E., Garcia-Casado, G., Witters, E., Inze, D., Long, J.A, De Jaeger, G., Solano, R, and Goossens, A (2010). NINJA connects the co-repressor TOPLESS to jasmonate signalling. Nature 464, 788–791.

Payne, C.T., Zhang, F., and Lloyd, AM. (2000). GL3 encodes a bHLH protein that regulates trichome development in arabidopsis through interaction with GL1 and TTG1. Genetics 156, 1349–1362.

Rerie, W.G., Feldmann, KA, and Marks, M.D. (1994). The GLABRA2 gene encodes a homeo domain protein required for normal trichome development in Arabidopsis. Genes & development 8, 1388–1399.

Sallets, A, Beyaert, M., Boutry, M., and Champagne, A. (2014). Comparative proteomics of short and tall glandular trichomes of Nicotiana tabacum reveals differential metabolic activities. Journal of proteome research 13, 3386–3396.

San-Bento, R, Farcot, E., Galletti, R, Creff, A, and Ingram, G. (2014). Epidermal identity is maintained by cell-cell communication via a universally active feedback loop in Arabidopsis thaliana. The Plant Journal 77, 46–58.

Schmidt, G.W., and Delaney, S.K (2010). Stable internal reference genes for normalization of real-time RT-PCR in tobacco (Nicotiana tabacum) during development and abiotic stress. Molecular genetics and genomics : MGG 283, 233–241.

Schnittger, A., Schöbinger, U., Stierhof, Y.-D., and Hülskamp, M. (2002). Ectopic B-Type Cyclin Expression Induces Mitotic Cycles in Endoreduplicating Arabidopsis Trichomes. Current Biology 12, 415–420.

Schnittger, A., Schöbinger, U., Stierhof, Y.-D., and Hülskamp, M. (2005). Ectopic B-Type Cyclin Expression Induces Mitotic Cycles in Endoreduplicating Arabidopsis Trichomes. Current Biology 15, 980.

Schnittger, A, Folkers, U., Schwab, B., Jürgens, G., and Hülskamp, M. (1999). Generation of a spacing pattern: the role of triptychon in trichome patterning in Arabidopsis. The Plant cell 11, 1105–1116.

Serna, L., and Martin, C. (2006). Trichomes: different regulatory networks lead to convergent structures. Trends in plant science 11, 274–280.

Shen, Q., Liu, Z., Song, F., Xie, Q., Hanley-Bowdoin, L., and Zhou, X. (2011). Tomato SlSnRK1 protein interacts with and phosphorylates βC1, a pathogenesis protein encoded by a geminivirus β-satellite. Plant physiology 157, 1394–1406.

Szemenyei, H., Hannon, M., and Long, J.A. (2008). TOPLESS Mediates Auxin-Dependent Transcriptional Repression During Arabidopsis Embryogenesis 319, 1384–1386.

Valkama, E., Salminen, J.P., Koricheva, J., and Pihlaja, K (2003). Comparative analysis of leaf trichome structure and composition of epicuticular flavonoids in Finnish birch species. Annals of Botany 91, 643–655.

Wada, T., Tachibana, T., Shimura, Y., and Okada, K. (1997). Epidermal cell differentiation in Arabidopsis determined by a Myb homolog, CPC. Science 277, 1113–1116.

Walker, AR, Davison, P.A., Bolognesi-Winfield, AC., James, C.M., Srinivasan, N., Blundell, T.L., Esch, J.J., Marks, M.D., and Gray, J.C. (1999). The TRANSPARENT TESTA GLABRA1 locus, which regulates trichome differentiation and anthocyanin biosynthesis in Arabidopsis, encodes a WD40 repeat protein. The Plant cell 11, 1337–1349.

Wang, S., Hubbard, L., Chang, Y., Guo, J., Schiefelbein, J., and Chen, J.G. (2008). Comprehensive analysis ofsingle-repeat R3 MYB proteins in epidermal cell patterning and their transcriptional regulation in Arabidopsis. BMC plant biology 8, 1–13.

Wester, K, Digiuni, S., Geier, F., Timmer, J., Fleck, C., and Hiilskamp, M. (2009). Functional diversity of R3 single-repeat genes in trichome development. Development 136, 1487–1496.

Williams, L., and Fletcher, J.C. (2005). Stem cell regulation in the Arabidopsis shoot apical meristem. Current opinion in plant biology 8, 582–586.

Wu, R, and Citovsky, V. (2017a). Adaptor proteins GIR1 and GIR2. II. Interaction with the co-repressor TOPLESS and promotion of histone deacetylation of target chromatin. Biochemical and biophysical research communications 488, 609–613.

Wu, R, and Citovsky, V. (2017b). Adaptor proteins GIR1 and GIR2. I. Interaction with the repressor GLABRA2 and regulation of root hair development. Biochemical and biophysical research communications 488, 547–553.

Yan, T., Chen, M., Shen, Q., Li, L, Fu, X., Pan, Q., Tang, Y., Shi, P., Lv, Z, Jiang, W., Ma, Y.N., Hao, X., Sun, X., and Tang, K (2016). HOMEODOMAIN PROTEIN 1 is required for jasmonate-mediated glandular trichome initiation in Artemisia annua. New Phytologist 213, 1145–1155.

Yang, C., Gao, Y., Gao, S., Yu, G., Xiong, C., Chang, J., Li, H., and Ye, Z. (2015). Transcriptome profile analysis of cell proliferation molecular processes during multicellular trichome formation induced by tomato Wo (v) gene in tobacco. BMC genomics 16, 868.

Yang, C., Li, H., Zhang, J., Luo, Z., Gong, P., Zhang, C., Li, J., Wang, T., Zhang, Y., and Lu, Y.E. (2011). A regulatory gene induces trichome formation and embryo lethality in tomato. Proceedings of the National Academy of Sciences of the United States of America 108, 11836–11841.

Yoo, S.D., Cho, Y.H., and Sheen, J. (2007). Arabidopsis mesophyll protoplasts: a versatile cell system for transient gene expression analysis. Nature protocols 2, 1565–1572.

