## Supplementary material for "NbCycB2 represses Nbwo activity via a negative feedback loop in the tobacco trichome developmemt": Fig.S1 - S11

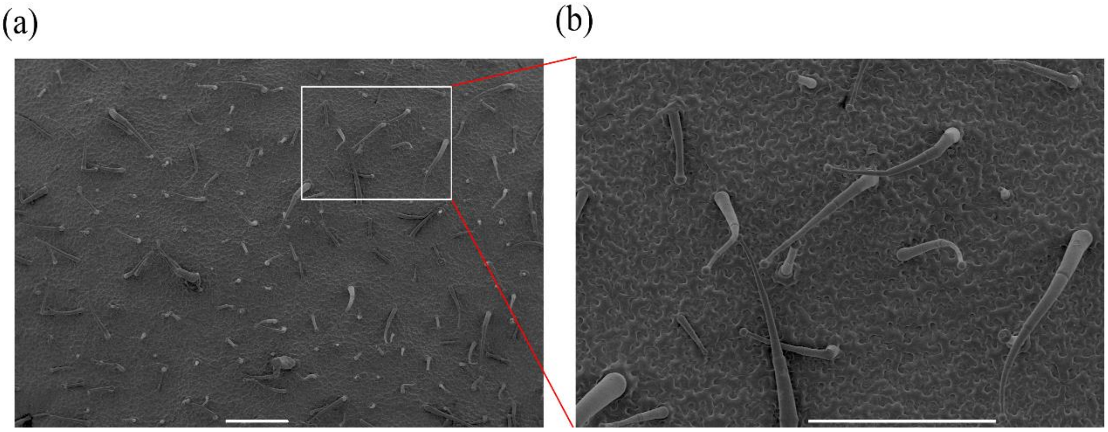


Fig. S1: The scanning electron micrographs (SEMs) of trichomes in the leaf of *N. benthamiana*

(a) The trichome types in the leaf of *N. benthamiana* were demonstrated. (b) The magnification of the white box area in figure a. The bar is 500 µm.

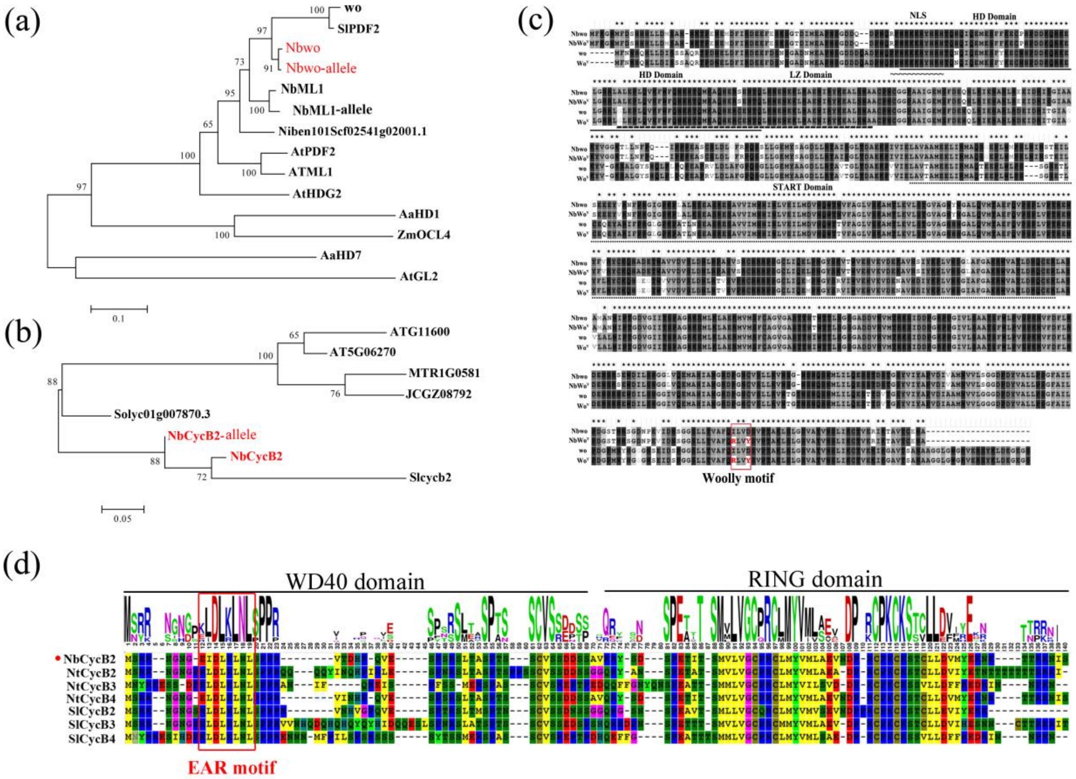


Fig. S2: Sequence analysis of Nbwo, NbCycB2 and their similar proteins

(a) Phylogenetic tree shown the relationships between Nbwo protein and other

HD-ZIP IV proteins. (b) Phylogenetic tree shown the relationships between NbCycB2 protein and other similar proteins. Protein sequence alignment and phylogenetic tree were performed in MEGA 5.2 software by using the maximum-likelihood (ML) criterion with 100 replicates bootstrap analysis. (c) Protein sequence alignment between Nbwo, NbWo$^V$, wo and Wo$^V$. Nbwo allele (NbWo$^V$) was mutant at 697 and 700 in the woolly motif [ isoleucine (I) changing to arginine (R) and aspartic acid (D) to tyrosine (Y)]. The red box is the Woolly motif sequences. (d) The analysis of NbCycB2 proteins conserved domains. The red box is the EAR motif protein sequences (KLDLKLNL).

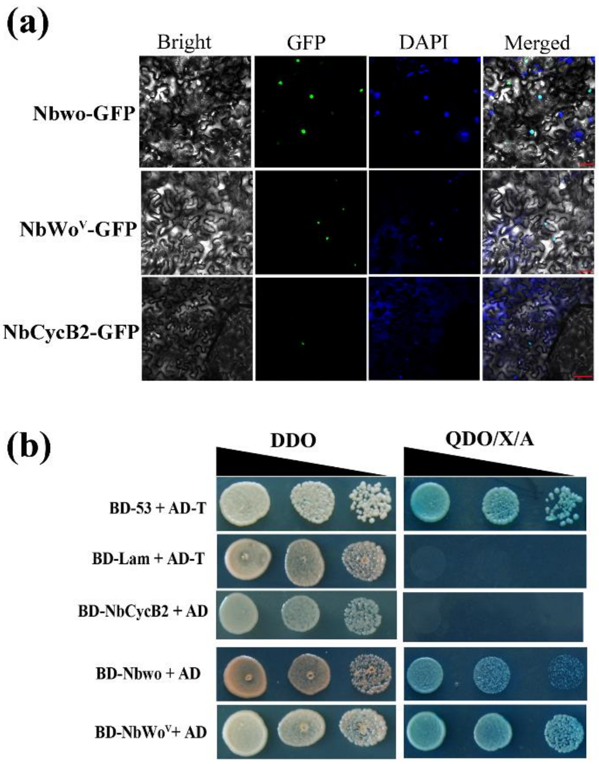


Fig. S3: The subcellular localization assay and auto activation test of *NbCycB2*, *Nbwo* and *NbWo$^V$*

(a) The subcellular localization of *NbCycB2*, *Nbwo* and *NbWo$^V$* were determined in *N. benthamiana* leaves. As shown in the fluorescence and bright field images, NbCycB2, Nbwo and NbWo$^V$ localized in the nucleus (bars, 50 μm).

(b) The auto activation test of *NbCycB2*, *Nbwo* and *NbWo$^V$* were shown. Blue clones grown on the QDO/X/A medium, which indicates these baits have autoactivation ability. Clones with pGADT7-53 (BD-53) and pGADT7-T (AD-T) as the positive controls, pGBKT7-Lam (BD-Lam) and pGADT7-T (AD-T) as the negative controls.

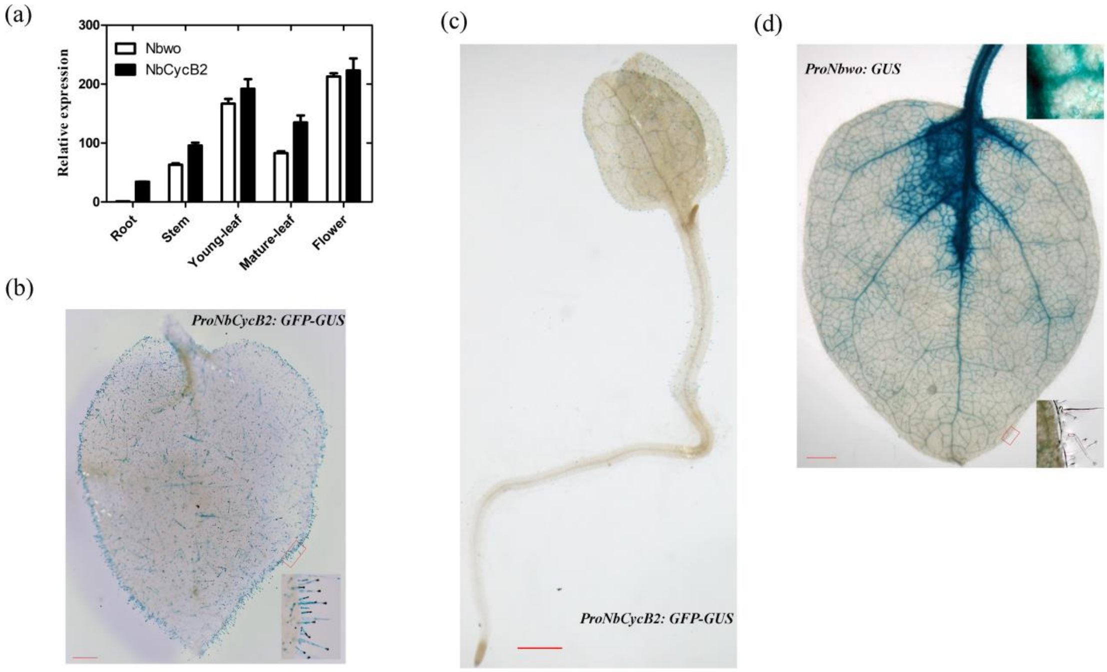


Fig. S4: The expression pattern of *NbCycB2* and *Nbwo* in *N. benthamiana*

(a) The relative expression levels of *NbCycB2* and *Nbwo* were measured respectively by real-time PCR in root, stem, young leaf, mature leaf and flower of *N. benthamiana*. Actin was used as internal reference, the expression of *Nbwo* in root as the control. Data are given as means SD (n = 3). (b), (c) The GUS staining were detected on the mature leaf and seedling of *ProNbCycB2*: *GFP-GUS* and *ProNbwo*: *GUS* transgenic lines (bars, 200 µm). (d) The GUS staining were detected on the young leaf of *ProNbwo*: *GUS* transgenic lines (bars, 200 µm).

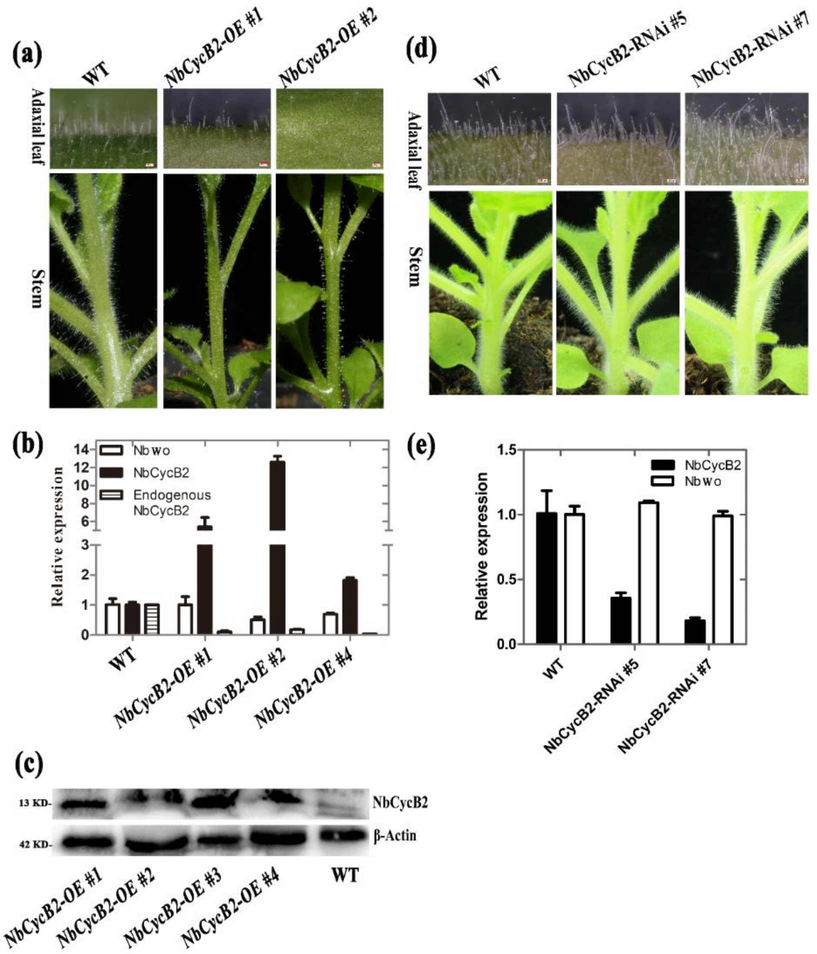


Fig. S5: Overexpression of *NbCycB2* and RNA interference of *NbCycB2* in *N. benthamiana*

(a) The trichome phenotypes on the stem and mature leaves of NbCycB2-OE (P35S: *NbCycB2*) transgenic lines were shown. The trichome density obviously reduced in the *NbCycB2-OE* lines. (b) The expression of *NbCycB2* and *Nbwo* were measured by real-time PCR in the *NbCycB2-OE* transgenic lines. Data are given as means SD (n = 3). (c) The Flag-NbCycB2 fused proteins were detected by using western blot in the *NbCycB2-OE* transgenic lines. (d) The trichomes derived from the stem and leaves of *NbCycB2-RNAi* transgenic lines. (e) The expression of exogenous *NbCycB2* and *Nbwo* were detected by real-time PCR in the *NbCycB2-RNAi* lines. Error bars represent SD (n = 3).

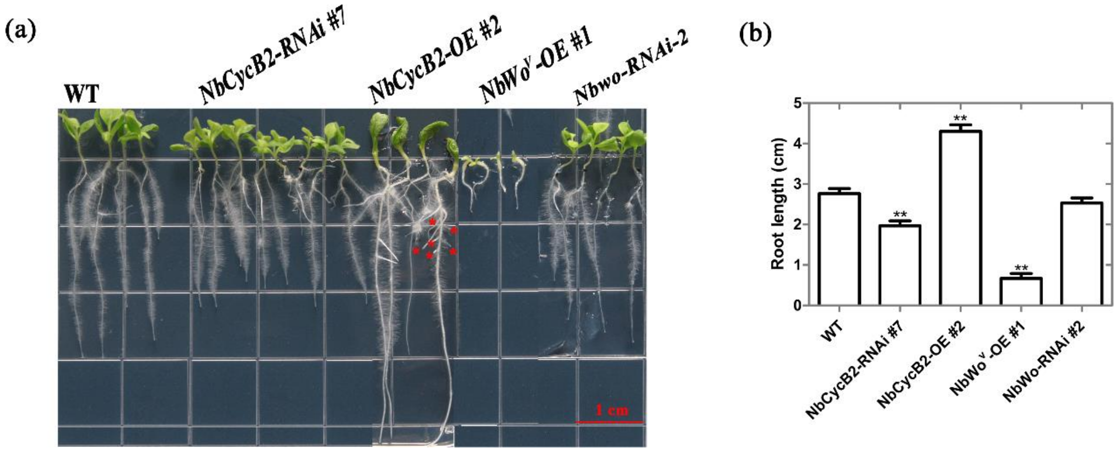


Fig. S6: The root phenotypes of wild type, *NbCycB2-RNAi* #7 T1, *NbCycB2-OE* #2 T1, *NbWo$^V$-OE* #1 T1, *Nbwo-RNAi #2* T1 seedlings

(a) The root phenotypes of wild type, *NbCycB2-RNAi* #7 T1, *NbCycB2-OE* #2 T1, *NbWo$^V$-OE* #1 T1, *Nbwo-RNAi #2* T1 two weeks-old-seedlings. The red bar is 1 cm. (b) The root length of wild type, *NbCycB2-RNAi* #7 T1, *NbCycB2-OE* #2 T1, *NbWo$^V$-OE* #1 T1, *Nbwo-RNAi #2* T1 two weeks-old-seedlings. "**" indicates a significant difference at P < 0.01 by Student's t test compared to WT. Error bars represent SD (n = 3).

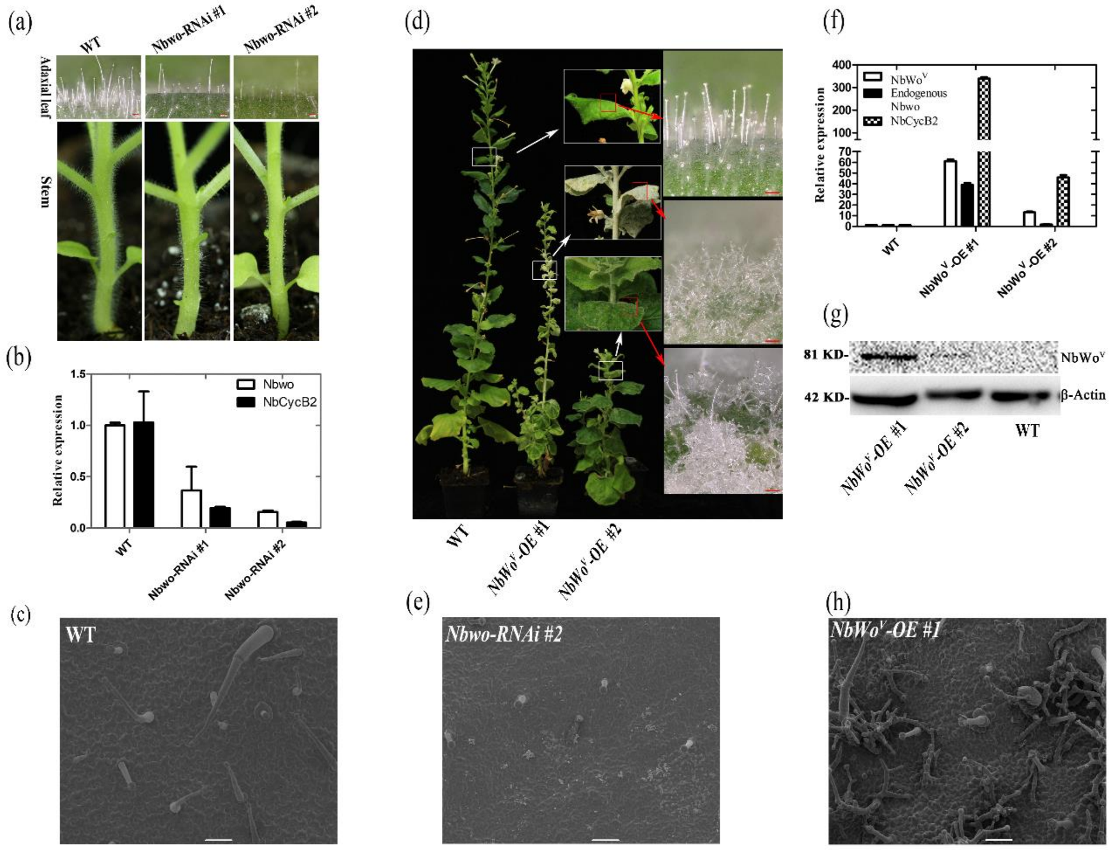

Fig. S7: RNA interference of *Nbwo* and overexpression of *NbWo*$^{V}$ in *N. benthamiana*

(a) The trichomes on stem and leaves derived from the *Nbwo-RNAi* transgenic lines. (b) The expressions of *NbCycB2* and *Nbwo* were measured by RT-PCR in the *Nbwo-RNAi* transgenic lines. Data are given as means SD (n = 3). (c), (e), (h) The SEMs of trichome in the mature leaves of wild type, *Nbwo-RNAi #2* and *NbWo*$^{V}$*-OE #1* lines was shown. The white bar is 100 µm. (d) The phenotype of *NbWo*$^{V}$*-OE* lines. Over expression of *NbWo*$^{V}$ induce clearly increase of trichomes density and branching. (f) The expression level of *NbWo*$^{V}$, *NbCycB2* and endogenous *Nbwo* were detected by RT-PCR in the *NbWo*$^{V}$*-OE* lines. Error bars represent SD (n = 3). (g) HA-NbWo$^{V}$ fusion protein detected by western blot in *NbWo*$^{V}$*-OE* transgenic lines.

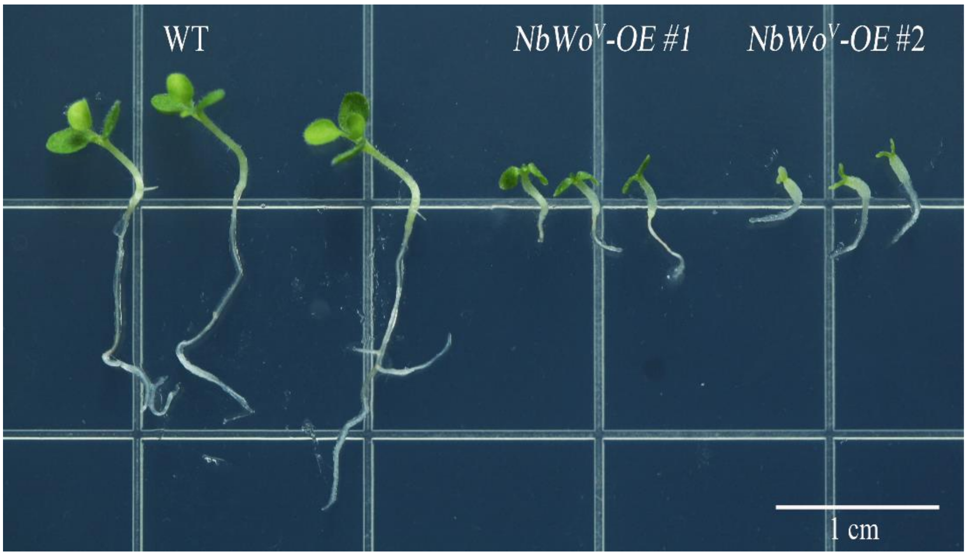


Fig. S8: The phenotype of *NbWo*$^{V}$*-OE* lines

Compare with WT, *NbWo*$^{V}$*-OE* #1*,* #2 lines shown a dwarfism (10-day-old seedlings).

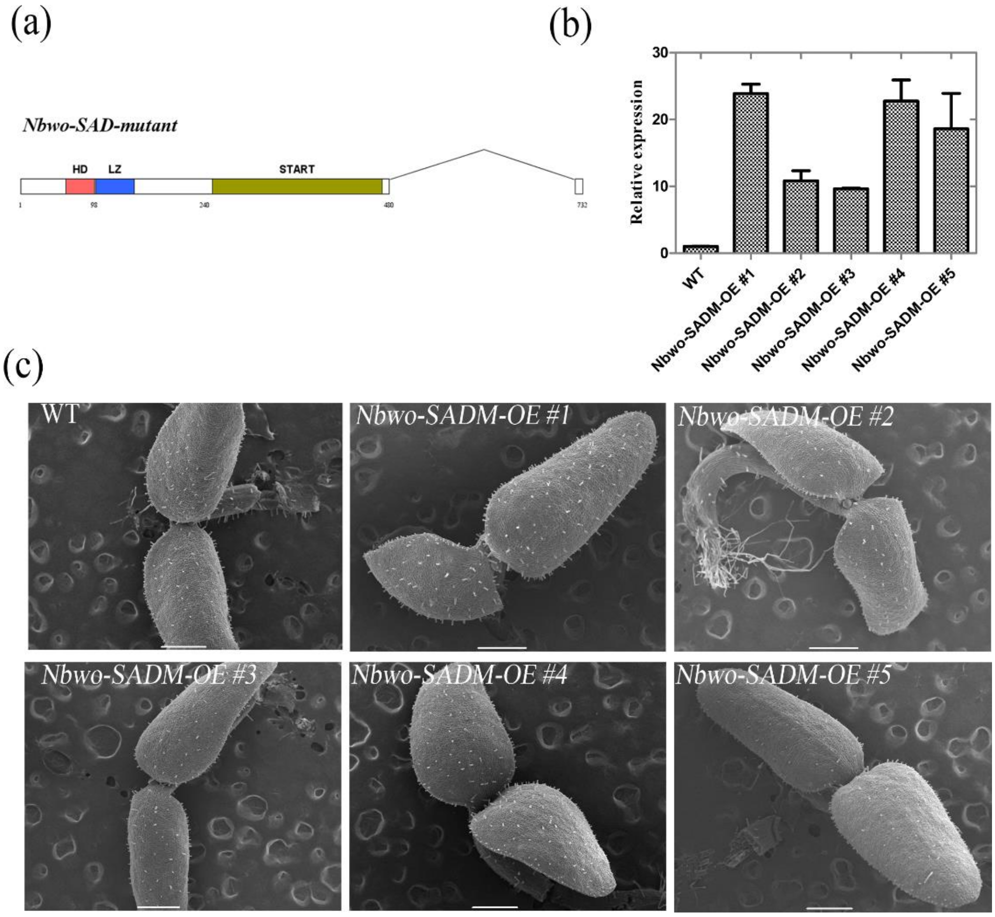


Fig. S9: The phenotype of overexpressing *Nbwo-SAD-mutant* in *N. benthamiana*.

(a) The Schematic diagrams of the mutantion of Nbwo protein SAD doman domain. (b) The relative expression levels of *Nbwo-SADM* were measured by qRT-PCR in F1 plants. Data are given as means SD (n = 3). (c) The phenotype of overexpression of *Nbwo-SAD-mutant* CDS (*Nbwo-SADM*) in the *N. benthamiana*. Compared with wild type, the trichome density was no significantly increase in the overexpression of *Nbwo-SAD-mutant* one week-old seedlings (*Nbwo-SADM-OE*). The white bar is 500 µm.

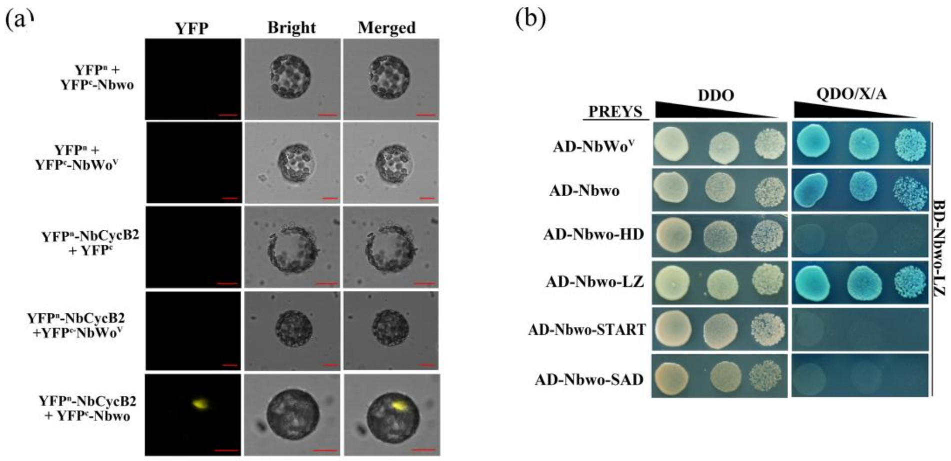


Fig. S10: the interaction between Nbwo and NbCycb2, Nbwo and Nbwo LZ domain.

(a) Interaction between NbCycB2 and Nbwo or NbWo$^V$ proteins were determined by BiFC assay in *N. benthamiana* protoplasts (bars, 20 μm). Yellow fluorescence indicates positive protein-protein interactions. (b) Detection of the interaction between the Nbwo and Nbwo LZ domain by using the yeast two-hybrid. Blue clones grown on the QDO/X/A medium indicates positive protein-protein interactions.

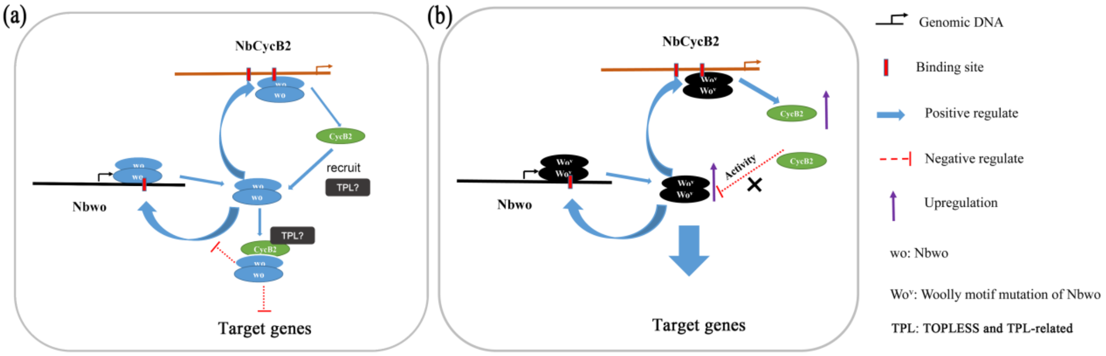


Fig. S11: A simplified model for regulation between *Nbwo* and *NbCycb2*

(a) Both Nbwo and NbWo$^V$ protein can dimerize itself into a homodimers, which then activate downstream genes to promote trichome initiation. Meanwhile, the homodimers also can promote the expression of *NbCycb2* and regulate itself endogenous expressing through bind to the *NbCycb2* promoter and itself genomic sequences respectively. In contrast, NbCycb2 binds to the Nbwo dimer and may inhibit the initiation of the trichome by inhibiting the transactivation of Nbwo. However, whether NbCycb2 participates in inhibitory activities through the recruitment of TOPLESS-like co-repressor requires further study. In summary, *Nbwo* and *NbCycb2* form a negative feedback loop to regulate the trichome development. (b) Since NbWo$^v$ does not interact with NbCycB2, it is protected from inhibition by NbCycB2 and promotes expression of its downstream genes, which results in a significant increase in the density and branch of the trichomes in the *NbWo$^v$* mutant lines.
